# GATA-regulated transcriptional program dictate cell fate equilibrium to establish the maternal-fetal exchange interface and fetal development

**DOI:** 10.1101/2023.06.19.545618

**Authors:** Ananya Ghosh, Rajnish Kumar, Ram Kumar, Soma Ray, Abhik Saha, Namrata Roy, Purbasa Dasgupta, Courtney Marsh, Soumen Paul

## Abstract

The placenta establishes a maternal–fetal exchange interface to transport nutrients and gases between the mother and the fetus. Establishment of this exchange interface relies on the development of multinucleated syncytiotrophoblasts (SynT) from trophoblast progenitors and defect in SynT development often leads to pregnancy failure and impaired embryonic development. Here, we show that mouse embryos with conditional deletion of GATA2 and GATA3 in labyrinth trophoblast progenitors have underdeveloped placenta and die ∼ embryonic day 9.5 (E9.5). Single cell RNA Seq (scRNA-Seq) analysis revealed excessive accumulation of multipotent labyrinth trophoblast progenitors upon conditional deletion of GATA factors. The GATA factor-deleted multipotent progenitors were unable to differentiate to matured SynTs. We also show that the GATA factor-mediated priming of trophoblast progenitors for SynT differentiation is a conserved event during human placentation. Loss of either GATA2 or GATA3 in cytotrophoblast (CTB)-derived human trophoblast stem cells (human TSCs) drastically inhibits SynT differentiation potential. Identification of GATA2 and GATA3 target genes along with comparative bioinformatics analyses revealed that GATA factors directly regulate hundreds of common genes in human TSCs, including genes that are essential for SynT development and implicated in preeclampsia and fetal growth retardation. Thus, our study uncovers a conserved molecular mechanism, in which coordinated function of GATA2 and GATA3 promote trophoblast progenitor-to-SynT commitment ensuring establishment of the maternal–fetal exchange interface.

## Introduction

Rodents and humans have hemochorial placentation, which is characterized by two vascular systems, maternal and fetal, and the maternal blood comes in direct contact with multinucleated SynTs (1–3). SynTs ensure the barrier function and facilitates nutrient and gas exchange between the maternal blood and the developing fetus (4). Establishment of this exchange interface is essential for the successful progression of pregnancy and mammalian reproduction.

In a post implantation mouse embryo mutipotent trophoblast stem and progenitor cells (TSPCs) arise within the ectoplacental cone (EPC) and the chorion (5). Eventually, the TSPCs within the EPC and chorion give rise to lineage-specific trophoblast precursors, which differentiate to trophoblasts cells of specialized functions of a matured placenta. The EPC and the chorion give rise to a functional placenta that comprises two structural regions-the avascular junctional zone (JZ) and the labyrinth zone (LZ), which contains SynTs and fetal vasculature. The JZ comprises of a compact layer of cells sandwiched between the labyrinth and the parietal trophoblast giant cell (p-TGC) layer. The TSPCs within the EPC contribute to four differentiated trophoblast cell types: TGCs (6), spongiotrophoblast cells (SpT) (7), glycogen cells (GC), and invasive trophoblast cells that invade the uterine wall and maternal vessels (8–10). The mouse placental labyrinth lies beneath the JZ and is characterized to maximize the exchange interface between the mother and the fetus (11).

The labyrinth development in the mouse placenta is initiated ∼E8.0 when the TSPCs of the chorionic ectoderm attaches with the allantoic mesoderm (12). At the onset of labyrinth formation, glial cells missing 1 (*Gcm1*) expression is induced in the TSPCs of the chorionic ectoderm (13). Post attachment, the chorio-allantoic layer starts invaginating at the sites of GCM1 expression (14), thereby creating branch point initiation. The labyrinth undergo extensive branching morphogenesis and expands from E9.5-E14.5 to establish a functional maternal-fetal interface with enormous surface area for nutrient and gas exchange (12, 13, 15). The maternal blood sinusoids and fetal blood capillaries within a mouse labyrinth are seperated by three differentiated trophoblast cell layers, the mononuclear sinusoidal trophoblast giant cells (S-TGCs), and two multinucleated Syncytiotrophoblasts, SynTI and SynTII (11, 16–19). S-TGCs, lining the maternal blood sinusoids of the labyrinth zone, express hormones and growth factors and are loosly attach to the SynTI layer (11, 12, 20). SynTI layer comes in direct contact with maternal blood due to the perforated nature of the S-TGC layer. The SynTII layer lies between the SynTI layer and the fetal endothelium. Thus, the three trophoblast layers along with the fetal endothelium form the maternal-fetal exchange-interface, which is integral for the barrier and exchange function of the placenta to assure embryonic growth and viability.

Studies on mutant mouse embryos have identified molecular regulators, such as peroxisome proliferator-activated receptor gamma (PPARG), GCM1, distal-less homeobox 3 (DLX3), CAAT enhancer binding protein alpha and beta (CEBPA/B), hepatocyte growth factor (HGF) and its receptor the Met tyrosine kinase (c-MET), which are important for matured syncytiotrophoblast development and placental labyrinth formation (14, 21–25). In addition, expression analyses identified markers that are specifically expressed in distinct trophoblast cell layers of the developing placental labyrinth. For example, mRNA expressions of nuclear receptor subfamily 6 group A member 1 (*Nr6a1),* ESX homeobox 1 *(Esx1)* along with *Gcm1, Dlx3, Cebpa/b* and *c-Met* can be specifically detected in the SynT cells during early stages of Labyrinth development (11). Furthermore, several trophoblast layer-specific markers for three labyrinth trophoblast cell-types have also been identified. For example, retroviral gene, *Syncytin a (Syna),* the Monocarboxylic acid transporter 1 (MCT1, encoded by *Slc16a1* gene) and transferrin receptor *(Tfrc)* are selectively expressed in SynTI, whereas, *Syncytin b (Synb),* MCT4 *(encoded by Slc16A3* gene) and iron transporter ferroportin, MTP1 (encoded by the gene *Slc40a1*), are selectively expressed in SynTII population (26–28). Similarly, the S-TGC cells selectively express Heart and neural crest derivatives expressed 1 (*Hand1)* and cathepsin Q (*Ctsq)* (6). The studies on mouse mutants provide significant insight about the structural architecture, molecular regulators and specific markers of the three trophoblast cell layers in the mouse labyrinth. However, we still have a poor understanding about the molecular processes through which they develop during placentation, especially for a long period of time it was unknown whether they arise from a common progenitor population during labyrinth development.

The presence of a common labyrinth progenitor in mouse was first reported by Ueno et al. (16) showing that mouse labyrinth contains an Epcam(hi) multipotent labyrinth trophoblast progenitor (LaTP), which can differentiate to all labyrinth trophoblast subtypes when cultured at a clonal level. Recent study by Marsh et al. (29), which utilized unbiased snRNA-seq followed by RNA-velocity mapping, further confirmed that LaTPs of a mouse placenta have trilineage potential to form-SynTI, SynTII and S-TGCs. The transition of LaTPs to the corresponding mature differentiated trophoblast populations relies on establishment of distinct gene expression patterns (29). The snRNA-Seq analyses showed that LaTPs that start expressing higher levels of *c-Met* get committed to the SynTII lineage and are identified as SynTII precursors. A similar observation had been previously reported showing that HGF/C-MET axis and WNT signaling mechanism are essential for the polarization and the terminal differentiation of the SynTII cells (30). LaTPs that express higher levels of *Egfr* and *Epcam* are biased towards forming the SynTI precursors, whereas the expression of *Lifr* and *Podxl* mark the transition and commitment of LaTPs to the S-TGC lineage. Interestingly, Natale *et al.*(*31*) reported existence of a SCA1 (encoded by *Ly6a* gene)-expressing multi-potent trophoblast progenitor population in the mid-gestation mouse placenta. These SCA1-expressing progenitors are reported to have potential to differentiate to trophoblast cell-types of both labyrinth zone and junctional zone. Collectively, these studies provide evidence that trophoblast development in mouse labyrinth is associated with establishment of multiple progenitor cell types that could adopt distinct differentiation fates based on gene expression programs. However, transcriptional mechanisms that control differentiation fates in these progenitors are still incompletely understood.

In humans, two types of SynT arise during different stages of placentation. At the site of blastocyst implantation, a primitive syncytium is formed (32–34) and erodes the surface of the uterine epithelium. This primitive syncytium lays the groundwork for the columns of cytotrophoblasts (CTBs) to penetrate the primitive syncytium and form the primary villi. Subsequently, proliferation and differentiation of the CTBs cause the primary villi to branch and mature into a villous placenta that comprises of two types of mature villi-the floating villi and the anchoring villi. The floating villi is bathed in maternal blood and comprises of an underlying layer of CTBs that eventually fuse to form a multinucleated outer layer of SynT (35, 36). Like in mice, the SynT layer in humans is multinucleated and comes in direct contact with maternal blood. Thus, formation of differentiated, multinucleated SynTs from their committed trophoblast progenitors for the establishment of the placental exchange surface is a conserved adaptation in both mice and humans(35, 37). Furthermore, several transcription factors, such as PPARG, GCM1 and DLX3, which are essential for placental labyrinth development in mice, have also been implicated in human SynT development (38–41). The CTB progenitors in anchoring villi, which anchors to the maternal decidua through CTB cell columns, adopt a different differentiation fate and develops into invasive extravillious trophoblast cells (EVTs) (42). Interestingly, recent single cell RNA-sequencing studies discovered existence of multiple CTB subpopulations, with distinct gene expression patterns, in a developing human placenta (43, 44). Thus, the presence of distinct CTB subpopulations and their altered differentiation fates in floating vs. anchoring villi indicate that the CTB to SynT and CTB to EVT transition during human placentation needs fine tuning of the gene expression program in CTB progenitors. However, transcriptional mechanisms that regulate gene expression programs in CTB progenitors to promote either SynT differentiation or EVT differentiation is incompletely understood.

Earlier, we and other laboratories showed that GATA family transcription factors GATA2 and GATA3 (henceforth mentioned as GATA factors) are conserved in the trophoblast cells of both mouse and humans placentae (45–49). Individual deletion of either *Gata2* or *Gata3* in mouse does not overtly affect plaecental development. *Gata3-*null mouse embryos die at ∼E11.5 due to defective neuroendocrine system development (50, 51) and *Gata2-*null mouse embryos die at ∼E10.5 due to defective hematopoiesis (52). However, combinatorial deletion of both *Gata2* and *Gata3* genes in an early post-implantation mouse embryo (GATA-DKO embryo) abrogates placentation leading to embryonic death (53) at around E 9.0. Despite this importance of GATA factors in early placentation, it was unknown whether cell-autonomous GATA factor function in labyrinth progenitors are essential for the SynT/labyrinth zone development and progression of pregnancy. Also, both GATA3 and GATA2 expressions are conserved in the human trophectoderm (54), which differentiates to primitive syncytium, as well as in CTB progenitors and in differentiated SynTs of human placentae. However, importance of GATA factors during human SynT development and their contribution in human pregnancy associated disorders is unknown.

In this study, we show conditional deletion of GATA factors in mouse labyrinth progenitors arrests them in a multipotent state leading to defective placental labyrinth development. The loss of GATA factors leads to defective lineage segregation and inability of the labyrinth progenitors to differentiate into mature SynTs. Similarly, loss-of-function of either GATA2 or GATA3 in human trophoblast stem cells (human TSCs) arrests them in an intermediate stage where they are poised for differentiation but are unable to undergo SynT differentiation. Thus, we elucidate a developmental stage-specific transcriptional program established by conserved GATA factors that prime differentiation of multipotent trophoblast progenitor to SynT lineage at the maternal-fetal exchange interface.

## Results

### GATA Factors function in mouse labyrinth trophoblast progenitors is essential for placentation and fetal development

As GATA factors are expressed extensively in the mouse labyrinth zone and their global loss abrogates labyrinth zone development (53), we wanted to investigate importance of GATA factors in a labyrinth progenitor-specific manner. To restrict GATA factor deletion to the labyrinth progenitors, we used a previously established *Tg(Gcm1-cre)1Chrn* (*Gcm1^Cre^*) mouse model (55), where the Cre recombinase expression is restricted to the *Gcm1* expressing cells. During mouse placentation, *Gcm1* is first expressed in trophoblast progenitors of the chorionic ectoderm at ∼E8.0 (13, 14). After chorionic attachment, the GCM1 expressing progenitors initiate the branching morphogenesis of the labyrinth and contribute to the development of both SynTI and SynTII cells (13, 14, 19, 29, 30).

To validate the specificity of the Cre expression, we crossed *Gcm1^Cre^*mice with *Gt(ROSA)26Sor^tm4(ACTB-tdTomato,-EGFP)Luo^/J*, (also known as mT/mG mice), which possess loxP flanked membrane-targeted tdTomato (mT) cassette and express strong red fluorescence in all tissues (56). Upon breeding with the *Gcm1^Cre^* recombinase expressing mice, the resulting progenies have the mT cassette deleted in the Cre expressing tissue(s), allowing expression of the membrane-targeted EGFP (mG) cassette located in-frame immediately downstream. Microscopic analyses of the conceptuses from cross between *Gcm1^Cre^* male and *mT/mG* female confirmed EGFP expression only in the labyrinth zone of the placentation site at ∼ E9.0 (Fig. 1A). These observation indicated the specificity of *Gcm1^Cre^* recombinase expression in the trophoblast cells of the labyrinth zone. Furthermore, at E12.5, EGFP expression is specifically noticed throughout the placental labyrinth (Supplementary Fig. S1A) indicating that that *Gcm1* expressing progenitors could contribute towards SynT populations throughout the developed labyrinth zone.

**Figure 1.**
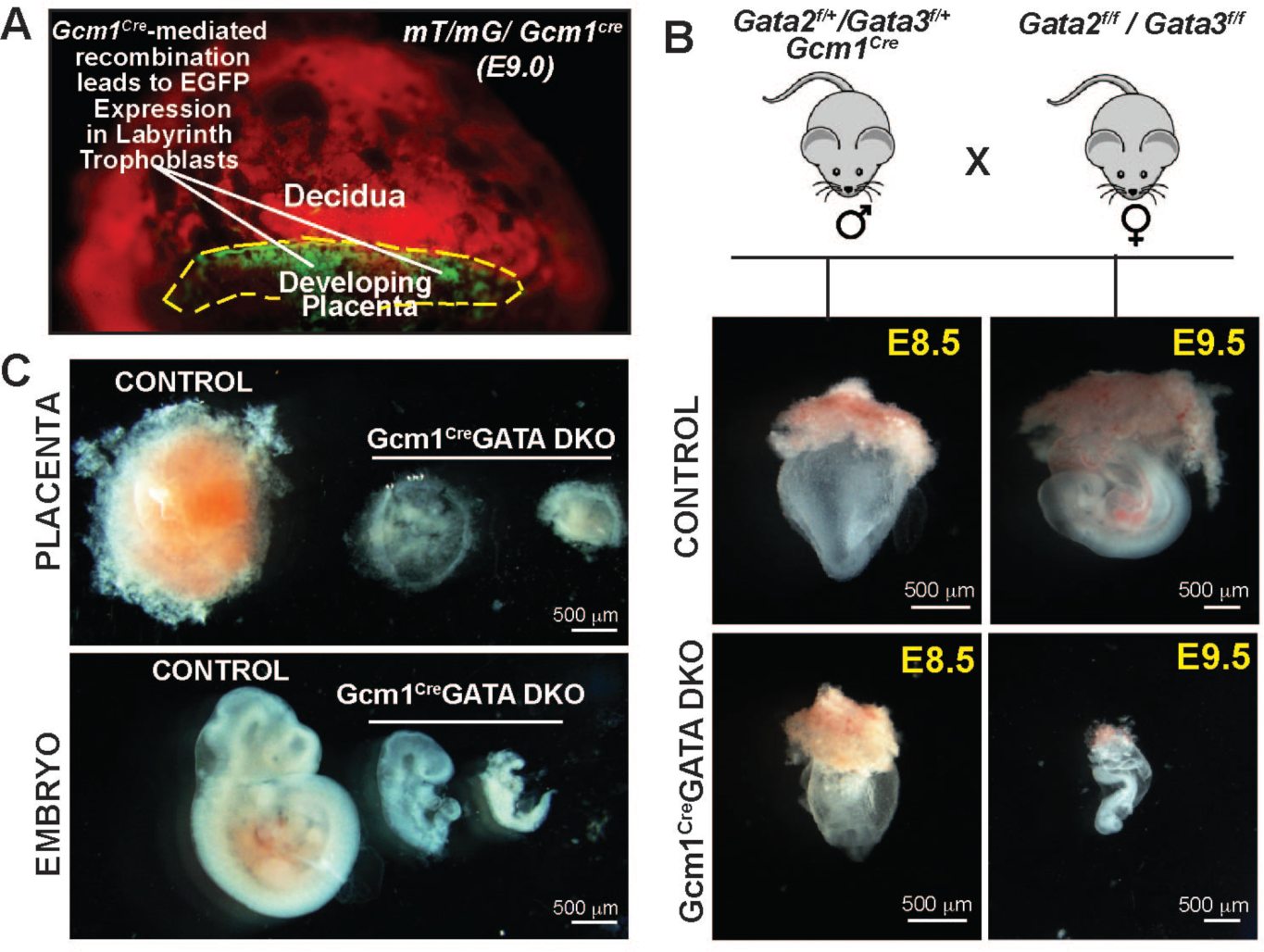
Conditional deletion of *Gata2* and *Gata3* factors in mouse labyrinth progenitors abrogate placental and embryonic development. **(A)** Validation of CRE recombinase expression specifically in labyrinth progenitors of a E9.5 *Gcm1^cre^*mT/mG conceptus. The CRE-mediated recombination of the mT/MG locus in the labyrinth proenitors lead to the green fluorescence. Green= Cre recombination; Red= no Cre recombination. **(B)**(top)Mating strategy used to determine the importance of *Gata2* and *Gata3* in labyrinth trophoblasts affecting placentation. Homozygous *Gata2^f/f^/Gata3^f/f^* female mice were crossed with heterozygous *Gata2^f/+^/ Gata3^f/+^/ Gcm1^Cre^* male mice to generate homozygous *Gcm1^Cre^*GATA DKO (*Gata2^f/f^/Gata3^f/f^/ Gcm1^Cre^*). Phenotype of mouse conceptuses derived using this mating strategy validates the importance of GATA2 and GATA3 in labyrinth trophoblasts affect placentation (*Gata2^f/f^/Gata3^f/f^* conceptuses were used as control). Embryonic and placental developments were analyzed at E8.5 and E9.5, representative images are shown. At E8.5, no obvious defect was observed in *Gcm1^Cre^*GATADKO embryos. However, defect in placentation and embryonic development was obvious in *Gcm1^Cre^*GATA DKO conceptuses at E9.5. (Scale bars, 500µm). **(C)** Representative images showcase developing placenta and embryos from control and *Gcm1^Cre^*GATA DKO. The *Gcm1^Cre^*GATA DKO placentae and embryos show varying severity in the phenotype depending on the penetrance of the Cre activity (Scale bars, 500µm.).

To delete GATA factors in labyrinth progenitors we studied mouse models, harboring conditional knockout alleles of either *Gata2* (*Gata2^f/f^)* or *Gata3* (*Gata3^f/f^)* or both of them (*Gata2^f/f^/Gata3^f/f^)* (53). To confine the homozygous deletion of *Gata2 and Gata3* gene only within the developing embryos, we crossed floxed female mice with male mice harboring *Gcm1^Cre^* allele along with respective floxed alleles. We noticed that individual loss of either *Gata2* or *Gata3* in labyrinth progenitors did not affect embryonic development as homozygous mutant pups were born with normal Mendelian ratio and without any noticeable defects. However, when we crossed *Gata2^f/f^/Gata3^f/f^* (henceforth mentioned as Control Floxed) female mice with male mice harboring *Gcm1^Cre^* allele and heterozygous floxed alleles of both *Gata2* and *Gata3* genes (*Gcm1^Cre^/Gata2^f/+^/Gata3^f/+^),* the embryos with the genotype *Gata2^f/f^*/*Gata3^f/f^/Gcm1^Cre^*(henceforth mentioned as *Gcm1^Cre^*GATA DKO) showed prominent developmental defect at E9.5 and were resorbed by E10.5 (Fig.1B, and Supplementary Fig. S1B, C and D). At E9.5, the *Gcm1^Cre^*GATA DKO placentae were significantly smaller in size in comparison to littermate controls (Fig.1C). The embryo proper of the *Gcm1^Cre^*GATA DKO embryos also showed arrested development (Fig.1C). Thus, our gene knockout studies confirmed that GATA factor functions in the labyrinth progenitors of a developing mouse embryo are essential for progression of placentation and embryonic survival beyond E9.5.

### Loss of GATA factors in the labyrinth progenitors impairs mature syncytia formation and disorganizes the trilaminar architecture at the maternal-fetal interface

The mouse maternal-fetal interface contains three layers of trophoblast cells, SynTI, SynTII and S-TGCs, that make up the trilaminar architecture (11, 17, 18). These layers facilitate the separation of maternal blood from the endothelial cells of the fetal vasculature. Thus, the two SynT layers along with the fetal endothelium are integral for the barrier and exchange function of the placenta to assure embryonic growth and viability. As we noticed defective placentation in *Gcm1^Cre^*GATA DKO conceptuses, we further analysed the labyrinth architecture in *Gcm1^Cre^*GATA DKO placentae.

Immunofluorescence analysis at E9.5 conceptuses revealed a significantly small labyrinth zone in *Gcm1^Cre^*GATA DKO placentae (Fig. 2A). Hence, we tested developmental status of the two mature SynT layers. We analyzed expressions of MCT1and MCT4, which are specifically expressed in SynTI and SynTII cell respectively (57). Unlike the control placentae, where tight juxtaposition of the two SynT layers was evident from MCT1 and MCT4 expressions, the *Gcm1^Cre^*GATA DKO placentae showed gross disruptions of both SynTI and SynTII layers (Fig. 2B and Supplementary Fig. S2A). The MCT4-expressing SynTII population were drastically reduced (Fig. 2B, and Supplementary Fig. S2A). In additon, strong reduction of MCT1-expressing SynTI population was also evident in *Gcm1^Cre^*GATA DKO placentae (Fig. 2B and Supplementary Fig. S2A). In contrast to SynT layers, development of p-TGCs, which border the placental-uterine interface, was not affected in *Gcm1^Cre^*GATA DKO placentae (Supplementary Fig. S2B)

**Figure 2.**
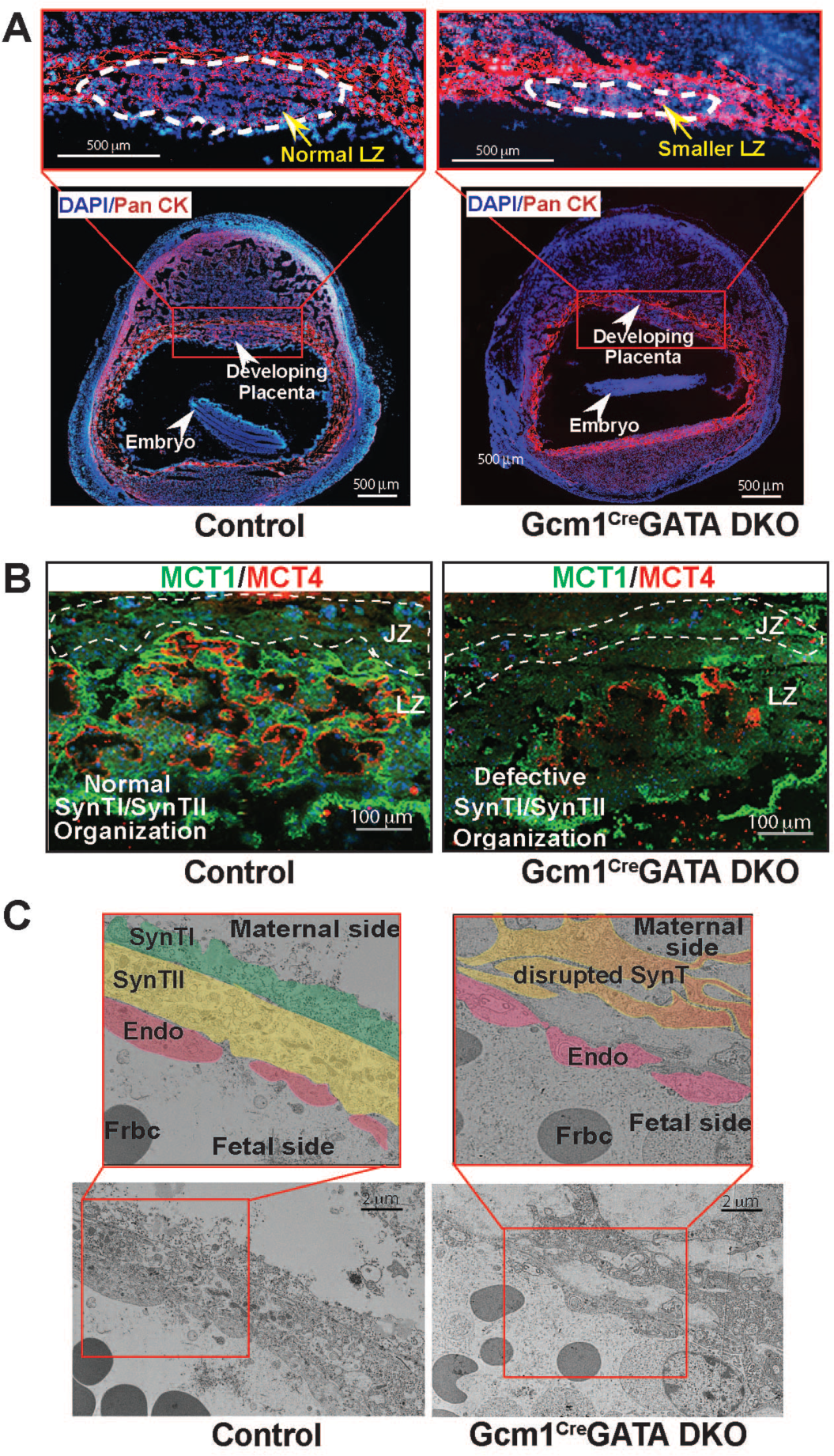
GATA factor function in labyrinth trophoblast progenitors is required for SynT maturation. **(A)** Placentation at control and *Gcm1^Cre^*GATA DKO implantation sites were analyzed at approximately E9.5 via immunostaining with anti–pan-cytokeratin antibody (red, trophoblast marker). The developing *Gcm1^Cre^*GATA DKO placenta lacks a well-developed labyrinth zone (white dashed line). Also, unlike in control embryos, the *Gcm1^Cre^*GATA DKO embryos are under-developed, and have a much smaller surface area. (Scale bars, 500 µm.). **(B)** Immunofluorescent analysis of the mature SynTI and SynTII layers was done in control and *Gcm1^Cre^*GATA DKO placental labyrinth with MCT1 (green, marks SynTI layer) and MCT4 (red, marks SynTII layer) antibodies. The developing *Gcm1^Cre^*GATA DKO placental labyrinth zone (LZ) defective mature SynTI and SynTI layers, they were disrupted as compared to nicely juxtaposed layered organization observed in the control placenta. (JZ, Junctional Zone, Scale bars, 100 µm). **(C)** The tri-laminar architecture of a developing labyrinth comprising of SynTI, SynTII and the fetal endothelium (endo), was captured using an electron microscope. In the control placenta, we observe the organized tri-layered architecture-SynTI (pseudo-color green), SynTII (pseudo-color yellow) and fetal endothelium (pseudo-color red) but in the *Gcm1^Cre^*GATA DKO both the SynT layers (pseudo-color orange) are completely disrupted. The endo layer in Gcm1Cre GATA DKO was not as severely affected. Indicating that the mature SynT layers do not form in Gcm1Cre GATA DKO placenta (Scale bars, 2 µm.). Frbc-fetal red blood cells, MCT1-Monocarboxylate transporter 1, MCT4-Monocarboxylate transporter 4, SynTI &SynTII-Syncytiotrophoblast layer I & II.

To further confirm the disruption of the labyrinth architecture we analyzed the knockout and control labyrinth under the electron microscope, which allowed us to visualize the trilaminar architecture at high resolution. Electron microscope analyses also revealed disrupted trilaminar structure in *Gcm1^Cre^*GATA DKO placentae (Fig. 2C). Instead of two closely positioned, mature SynT layers, the knockout labyrinth contained only one SynT layer that was discontinuous. Thus, from our immunofluorescence and EM studies, we concluded that the labyrinth progenitor-specific loss of GATA factors leads to defective development of the mature SynT lineages, resulting in abrogation of placental labyrinth formation.

### Single cell RNA Seq (scRNA-Seq) revealed the importance of GATA factors in committing the labyrinth progenitor to the SynT lineage

As loss of GATA factors impaired formation of mature SynT cells, we next asked whether the distribution of trophoblast progenitors are altered in *Gcm1^Cre^*GATA DKO placentae. As labyrinth progenitors are single cell populations, we performed single cell RNA Sequencing (scRNA-Seq) to capture distribution of distinct progenitor populations with E9.5 control and *Gcm1^Cre^*GATA DKO placentae. Mouse placenta from E9.5 embryos were carefully isolated and the embryo proper and decidual tissue were dissected out (Supplementary Fig. S3A). The embryo was used for genotyping to confirm *Gcm1^Cre^*GATA DKO placentae. The pooled control and *Gcm1^Cre^*GATA DKO placental tissues were used to generate single cell suspensions and subjected for scRNA-seq analyses.

Dimensionality reduction and clustering of all single cells was done using Seurat package and projected using t-SNE plots showcasing visible proximity between clusters. The Seurat package identified 21 clusters (Supplementary Fig. S3B). Based on the gene expression profile and cell type specific markers (mentioned in SI Appendix, Supplememental table 1) these clusters were classified into 5 broad groups: endodermal, endothelial, stromal, trophoblast and blood cells. Clusters 2, 4, 13, 14 were identified as trophoblast cells as they highly expressed trophoblast markers *Epcam, Ly6a, Prl3d1, Prl2c2, Tpbpa* (Supplementary Fig. S3B).

For a better understanding of the status of the trophoblast progenitors, we re-annotated only trophoblast cells using the Seurat package and identified eight distinct cell clusters based on marker gene expressions (Fig 3A and SI Appendix, Supplememental table 1). We observed a cell population (cluster 3, Fig. 3A) with high level expression of *Ly6a* (Fig 3B, C). These cells also highly express several other labyrinth progenitor-specific genes, such as *Wnt4,* T-Box transcription factor 3 *(Tbx3), Cebpb,* imprinted gene Iodothyronine Deiodinase 3 *(Dio3), and Tfrc* (Fig 3B, Supplementary Fig. S4A). Interestingly, the *Ly6a* expressing cell population also showed high level expressions of Epidermal Growth Factor Receptor (*Egfr)* (Fig 3C), which were earlier identified to be expressed in multipotent LaTPs by Marsh *et al.*(29). Thus, we concluded that the the *Ly6a, Cebpb, and Egfr* expressing cells comprises the multipotent LaTP compartment within the E9.5 labyrinth. We also identified more committed SynTI (cluster 6, Fig. 3A), SynTII (Cluster 5, Fig. 3A) and S-TGC (Cluster 8, Fig. 3A) precursors, through which these multipotent progenitors transition to the corresponding mature differentiated populations. The committed SynTI precursors showed high expressions of characteristic SynTI markers, *Tfrc* and *Slc16a1* (MCT1) (Fig. 3B,C, Supplementary Fig. S4A). *Nr6a1* and *Dlx3,* which are expressed in SynT cells during early labyrinth development (11) expressions, were detected in both SynTI and SynTII precursors (Supplementary Fig. S4B). The SynTII precursors highly express *Slc40A1* (MTP1), *Esx1, Fabp3, Phlda2, Wnt7b* and *Slc16a3* (MCT4) (Fig. 3B, C, Supplementary Fig. S4C). In addition, The S-TGC precursors express *Lifr* and *Podxl* (Fig. 3B, C, Supplementary Fig. S4D), which were earlier reported to be induced during S-TGC commitment (19). In addition, S-TGC precursors also highly express *Hand1,* which was mostly suppressed in multipotent labyrinth progenitors and in SynT precursors (Fig. 3B, C). Interestingly, similar to the snRNA-seq analyses by Marsh et al. (29), our scRNA-seq analyses detected high level of *Met* expression only in SynTII precursors (Supplementary Fig. S4C). The single cell trophoblast populations also contained four junctional zone progenitor trophoblasts (JZP1-4, Fig. 3A). JZP1-3 showed high level *Hand1* expression along with placenta-specific gene Placenta Enriched 1 (*Plac1*) (Fig. 3B,C). Whereas, JZP4 cells showed high level expression of SpTs marker, *Tpbpa,* along with other JZ trophoblast-specific genes *Prl8a9, Plac1, Pcdh12 and Ncam1* (Fig. 3A,B, Supplementary Fig. S4E).

**Figure 3.**
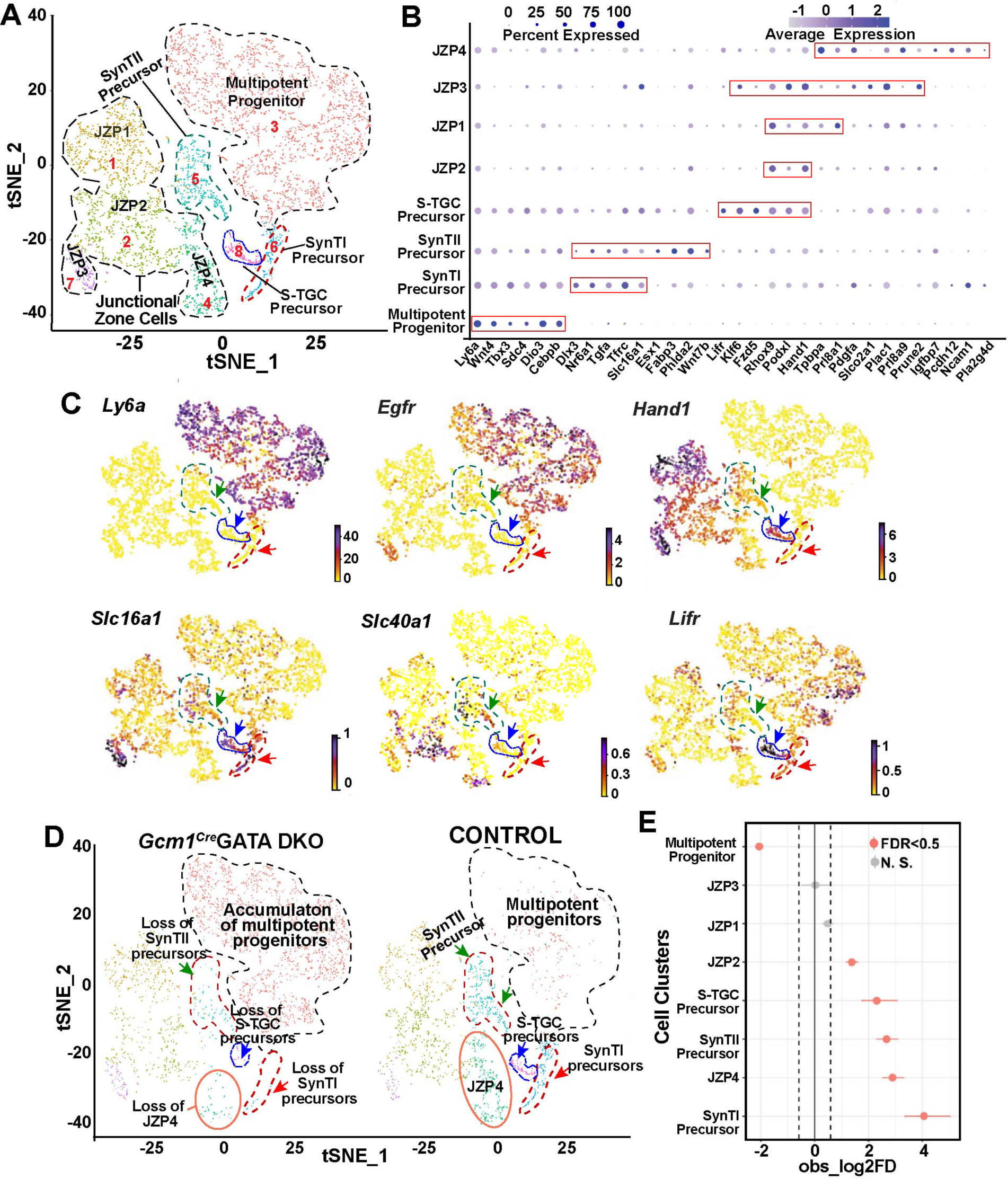
Loss of GATA factors impairs differentiation of multipotent labyrinth trophoblast progenitors to more committed precursors. **(A)** Visualization of the scRNA-Seq data with the analysis plotted in two dimensions by transcriptome similarity using t-distributed stochastic neighbor embedding (t-SNE). t-SNE plot represents the cells included in the trophoblast subset, clustered, and plotted according to transcriptome similarity. Clusters were annotated according to canonical marker genes. Each dot represents one cell colored according to assignment by clustering analysis (see Materials and methods). Dotted lines encircle clusters with common properties segregated into five trophoblast sub-clusters-Multipotent progenitors, SynTI precursors, SynTII precursors, S-TGC precursors and junctional zone cells. **(B)** Dot plot showing average expression and percent of cells in each cluster expressing canonical and novel marker genes identified for each cluster. Genes listed on the x-axis and clusters on the y-axis. **(C)** RNA velocity plots showing the expression of marker genes in specific clusters, the scale represents the level of expression-yellow (low), red (intermediate) and blue (high). The colored arrows mark specific populations-green=SynTII precursors, blue= S-TGC precursors and red= SynTI precursors. **(D)** Split by condition, t-SNE plot representing the difference in the trophoblast sub-clusters between control and Gcm1Cre GATA DKO samples. The t-SNE plot shows accumulation of the multipotent progenitors (cluster marked with black dotted lines) but loss of SynTI precursors, SynTII precursors and S-TGC precursor (clusters marked by red, green and blue arrows respectively), in the Gcm1Cre GATA DKO compared to the control. JZP3 cluster (marked in orange solid line) also diminishes in the Gcm1Cre GATA DKO placenta. **(E)** Single cell proportion test R-function was performed to generate a plot that highlights the difference between proportions of cells in each cluster between the control and Gcm1^Cre^ GATA DKO placental samples. The plot shows that cluster multipotent progenitor significantly increases in proportion in the Gcm1 Cre GATA DKO sample whereas SynTI and SynTII precursors, S-TGC pre-cursors, JZP2 and JZP3 show significant increase in proportion in the control sample. ZP1 and JZP4 do not show significant change in either of the samples.

The comparison of trophoblast progenitor clusters between control and *Gcm1^Cre^* GATA DKO placentae revealed two obvious differences. Compared to the control placenate, the *Gcm1^Cre^* GATA DKO placentae contained significantly higher number of multipotent progentior populations (Fig. 3D, E). In contrast, the committed SynTI, SynTII and S-TGC precursors, which express high levels of *Epcam* as reported earlier by Ueno et al. (16) (Supplementary Fig. S5A), were strongly diminished in the *Gcm1^Cre^* GATA DKO placentae (Fig. 3D, E, Supplementary Fig. S5A). Among the JZPs, JZP1 and JZP3 populations were not significantly altered in *Gcm1^Cre^* GATA DKO placentae, whereas cluster comprising JZP2 cells was slightly reduced. However, the JZP4 progenitors, which express *Tpbpa,* were also reduced in *Gcm1^Cre^* GATA DKO placentae (Fig. 3D, E, Supplementary Fig. S5B ).

To validate the scRNA-Seq observation we performed additional experiments. First, we confirmed enrichment of the *Ly6a* expressing multipotent progenitor population in the absence of GATA factors via fluorescence activated cell sorting (FACS) analysis. We generated single cell prepartion from control and *Gcm1^Cre^* GATA-KO placentae, removed all the hematopoietic and endothelial cells, and tested percentage of cells that were SCA1 (encoded by *Ly6a*) positive. We observed a significantly higher number of trophoblast cells that were SCA1 positive in the knockout placentae (Supplementary Fig. S6A). Next, we performed quantitative RT-PCR analyses to test expression of marker genes for SynTI, SynTII, S-TGC precursors and junctional zone trophoblasts in control and *Gcm1^Cre^* GATA DKO placentae. The RT-PCR analyses confirmed strong loss of *Nr6a1, Slc16a1,Fabp3, c-Met and Dlx3* in E9.5 *Gcm1^Cre^*GATA DKO placentae (Supplementary Fig. S6B). In contrast, mRNA expressions of *Podxl* (S-TGC precursor marker), *Hand1, Prl3d1* and *Prl2c2* (JZ trophoblast markeres) were not significantly altered in *Gcm1^Cre^* GATA DKO (Supplementary Fig. S6B). However, RT-PCR analyses also confirmed loss of *Tpbpa* expression in E9.5 *Gcm1^Cre^* GATA DKO placentae. Thus, our scRNA-seq, FACS and RT-PCR analyses indicated that the loss of the GATA factors in the labyrinth trophoblast progenitors arrest their development at a *Ly6a/Egfr* expressing uncommitted progenitor state, thereby affecting differentiation to committed SynTI, SynTII and S-TGC precursors. The strong loss of *Tpbpa*-expressing JZPs in *Gcm1^Cre^* GATA DKO placentae is an interesting observation and indicate that SpT development during mouse placentation might rely on crosstalk with labyrinth zone trophoblast precursors.

### Loss of GATA2 or GATA3 in human TSCs impairs SynT differentiation

The human placenta is comprised of chorionic villi that stem from the chorionic plate. These villous-like structures may anchor onto the uterine decidua (anchoring villi) or remain floating, bathed in maternal blood (floating villi). Floating villi have a characteristic cellular bilayer consisting of two types of trophoblast cells-an inner layer of mononuclear proliferative CTBs and an outer layer of multinucleated, differentiated syncytiotrophoblasts (SynT) formed by the fusion of underlying CTBs. In a matured term placenta, the CTB population is diminished and placental villi mainly contain differentiated SynT population. Our protein expression analyses revealed that both GATA2 and GATA3 are expressed in the CTBs and SynTs of a first-trimester human placenta, and their expression is maintained in SynTs within matured term placentae (Fig. 4A and Supplementary Fig S7A).

**Figure 4.**
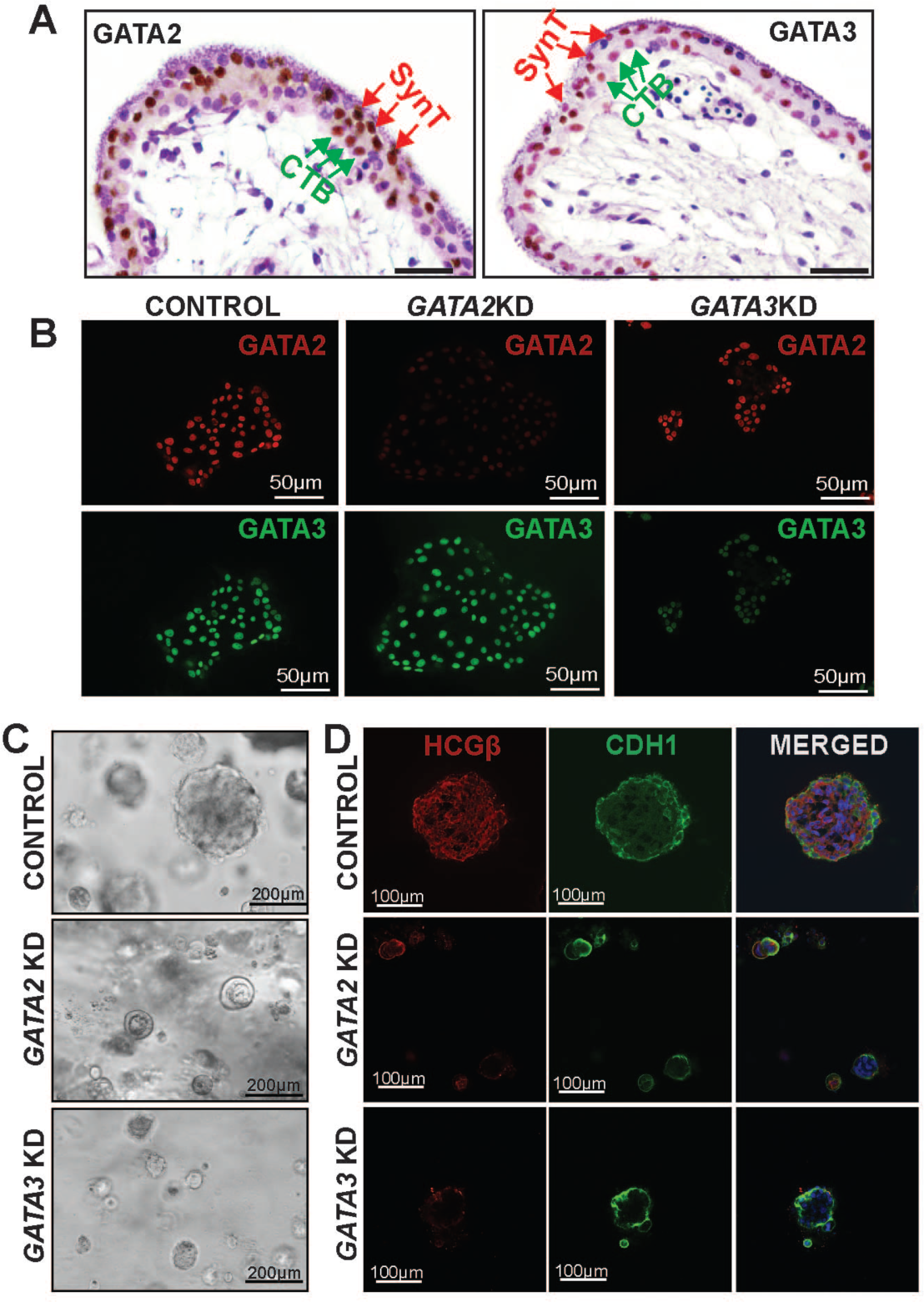
GATA2 and GATA3 optimizes self-renewal ability of human TSCs. **(A)** Immunohistochemistry showing expression of GATA2 and GATA3 in first trimester (9wk) human placenta. GATA2 (right) and GATA3 (left) are expressed abundantly in the syncytiotrophoblast cells (SynT,red arrows, outer layer) as well as within the villous cytotrophoblast progenitors (vCTBs, green arrows, inner layer) within the floating villi. (Scale bars, 50 µm.). **(B)** Immunofluorescence analysis confirming specific depletion of GATA2 and GATA3 protein expressions after shRNA-mediated knockdown in human TSCs. **(C)** 3D organoid culture was done with control, GATA2 KD and GATA3 KD human TSCs. The loss of function of GATA2 or GATA3 affected large organoid formations as observed in the control at day 8. The organoids formed with GATA2 KD and GATA3 KD human TSCs were much smaller in size and number. **(D)** Immunofluorescent analysis of the organoid integrity was done by highlighting its ‘inside-out’ model, with CTBs marked by E-Cadherin antibody (green) on the outside and SynT marked by hCGβ antibody (red) on the inside. The control organoids showcased a distinct inside-out model. The GATA2 KD and GATA3 KD organoids were smaller and underdeveloped, they lacked hCGβ (red) on the inside, indicating no SynT formation but had E-Cadherin (green) expression suggesting that the CTBs were not affected.

Recent studies showed that bonafide human TSCs, which were derived from first-trimester CTBs (58), are an excellent model to study molecular mechanisms that regulate CTB to SynT differentiation. Thus, we performed loss-of-function studies with human TSCs to test the importance of GATA2 and GATA3 in SynT development during human placentation. We used lentivirus-mediated delivery of short hairpin RNA (shRNAs) to specifically deplete *GATA2* and *GATA3* in human TSCs (Fig. 4B and Supplementary Fig. S7B). Interestingly, when cultured in 2-dimensional stem state culture conditions (2D stem-state culture, (58)) that maintain proliferating cultures, human TSCs with knockdown of either *GATA2* (*GATA2*KD human TSC) or *GATA3* (*GATA3*KD human TSCs) maintained the stem state colony morphology (Supplementary Fig. S7C) and did not show significant reduction in cell proliferation ability. However, combinatorial loss of both GATA factors in human TSCs (Dual *GATA*KD human TSC) impaired stem-state morphology and cell proliferation in a 2D stem-state culture condition (Supplementary Fig. S7C).

Interestingly, RT-PCR analyses showed that mRNA transcription factor *TEAD4*, cofactor vestigial like family member 1 (*VGLL1)*, and tumor protein P63 (*TP63),* which are critical to maintain human TSC stem state (37, 43, 59) as well as cell proliferation regulators polo like kinase 1 and 2 (*PLK1* and *PLK2)* and Wnt signaling component, *WNT5A,* were downregulated in *GATA2*KD and *GATA3*KD human TSCs (Supplementary Fig. S7D). Therefore, we also tested self-renewal ability of *GATA2*KD and *GATA3*KD human TSCs by assessing their ability to form self-renewing 3-dimensional (3D) trophoblast stem cell organoids (3D TSC organoid culture). In contrast to 2D-stem state culture, the 3D organoid cultures of both *GATA2*KD and *GATA3*KD human TSCs showed strong impairment in organoid formation ability (Fig. 4C). Control human TSCs formed large organoids with prolonged culture (7-10 days) and could be dissociated and reorganized to form secondary organoids, indicating the self-renewing ability. In contrast, *GATA2*KD and *GATA3*KD human TSCs formed much smaller organoids (Fig. 4C), which were not maintained upon passaging. Thus, we concluded that individual functions of GATA2 and GATA3 are important to maintain optimum self-renewal ability in human TSCs, especially when they are cultured in 3D culture condition. However, our findings in 2D-stem state culture also imply that GATA2 and GATA3 might have complementary functions in human TSCs that allow sustained proliferation and stem-state maintenance when any one of them are depleted in 2D-stem state culture. Whereas, irrespective of culture conditions, combinatorial loss of GATA2 and GATA3 abrogates stem-state maintenance in human TSCs. As human TSCs largely recapitulate the properties of CTB progenitors of a first-trimester human placenta, our findings strongly indicate that GATA factors are important to maintain the self-renewal ability in CTBs of a developing human placenta.

The 3D TSC organoids grow in an inside-out pattern. The undifferentiated TSCs in the outer layer undergo proliferation and express E-Cadherin (CDH1), whereas the inside cells undergo SynT differentiation and highly express HCGβ. We noticed that the defective organoid formations of *GATA2*KD and *GATA3*KD human TSCs are associated with near complete loss of HCGβ-expressing SynTs inside those organoids (Fig. 4D). Thus, our 3D TSC organoid culture strongly suggested that loss of either GATA2 or GATA3 in human TSCs might lead to defective SynT development. To confirm this aspect, we tested multinucleated SynT formation ability of *GATA2*KD and *GATA3*KD human TSCs in culture conditions that specifically promote SynT differentiation.

Human TSCs, when cultured in the presence of cAMP agonist forskolin, synchronously differentiate and fuse to form two-dimensional (2D) syncytia on a high attachment culture plate or 3D cyst-like structures on a low adhesion culture plate. In both 2D and 3D culture conditions, differentiated human TSCs highly express SynT markers such as HCGβ. We noticed that the loss of GATA2 in human TSCs abrogated SynT maturation when cultured in 2D-SynT culture conditions (Fig. 5A). *GATA2*KD cells failed to fuse and gradually became unhealthy. The loss of GATA3 in human TSCs also impaired SynT maturation in 2D-SynT culture conditions. *GATA3*KD cells retained their stem state like colony morphology and did not fuse to form multinucleated mature SynTs (Fig 5A). Impaired SynT differentiation was also confirmed by monitoring HCGβ and CDH1 expression. Unlike in control human TSCs, induction of HCGβ protein expression was strongly inhibited in *GATA2*KD and *GATA3*KD human TSCs (Fig. 5B), whereas expression of CDH1 was maintained along with a near-complete inhibition of cell-to-cell fusion (Fig. 5B).

**Figure 5.**
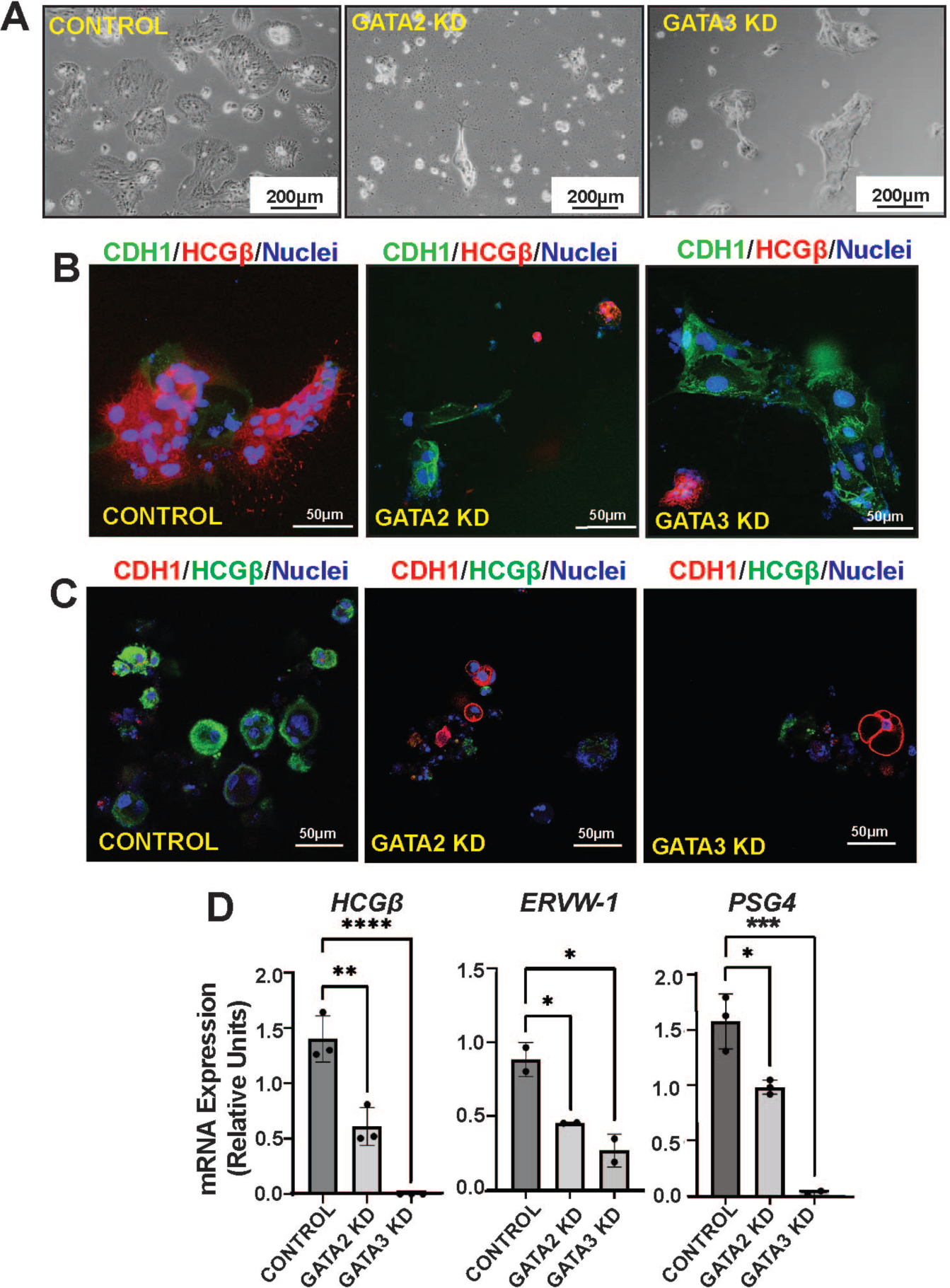
GATA2 and GATA3 are essential regulators for SynT development in human. **(A)** Control, GATA2 KD and GATA3 KD human TSCs were subjected to two-dimensional (2D) SynT differentiation on collagen-coated adherent cell culture dishes. Image panels show impaired ST(2D) colony formation in GATA2 KD and GATA3 KD cells. (Scale bars, 200 µm.). **(B)** Immunofluorescence analysis of the ST(2D) colonies (left) marked by the loss of E-Cadherin (CDH1, green) expression but induction of HCGβ expression (red). GATA3 KD ST(2D) show altered cellular morphology, maintenance of E-Cadherin (CDH1, green) expression and impaired induction of HCGβ expression (right). GATA2 KD ST(2D) colonies (middle) were much smaller, there was impaired cell fusion as E-Cadherin was maintained compared to the control but lacked HCGβ expression. **(C)** Immunofluorescence analysis of the ST(3D) colonies, similar to the 2D culture is marked by the loss of E-Cadherin (CDH1, red) expression but induction of HCGβ expression (green). Both GATA2KD and GATA3 KD ST(3D) show altered cellular morphology, maintenance of CDH1 expression and impaired induction of HCGβ expression. **(D)** Quantitative RT-PCR analyses (mean ± SE; n = 3, P ≤ 0.001) reveal impaired induction of SynT markers like CGβ, ERVW-1 and PSG4 in GATA2 KD and GATA3 KD human TSCs, undergoing 3D SynT differentiation.

Loss of SynT differentiation potential of *GATA2*KD and *GATA3*KD human TCSs was also evident under the 3D-SynT culture condition. Control human TSCs spontaneously formed larger sphere-like structures while the *GATA2*KD and *GATA3*KD human TSCs were unable to form larger spheres. Instead, they mainly formed smaller aggregates (Fig. 5C). Furthermore, like in 2D-SynT culture, we confirmed loss of HCGβ protein expression and retainment of CDH1 expression in 3D-SynT cultures of *GATA2*KD and *GATA3*KD human TSCs (Fig. 5C). mRNA expression analyses indicated significant inhibition of SynT-specific markers syndecan 1 (*SDC1), CGB*, endogenous retrovirus group W member 1 (*ERVW1)*, and pregnancy specific beta-1-Glycoprotein 4 (*PSG4)* mRNA inductions in the small cell aggregates formed by *GATA2*KD or *GATA3*KD human TSCs (Fig. 5D). Thus, both 2D and 3D-SynT differentiation systems revealed impaired SynT differentiation potential of the *GATA2*KD and *GATA3*KD human TSCs.

### GATA factors directly regulate expression of common genes that are important for SynT differentiation and are implicated in pathological pregnancies

To further understand how GATA2 and GATA3 functions promote SynT development in humans, we performed unbiased genomics studies. We performed RNA-Seq analysis with control, *GATA2*KD, and *GATA3*KD human TSCs when they were subjected to SynT differentiation to identify differentially expressed genes (DEGs) upon GATA2 and GATA3 depletion. RNA-seq analyses identified 4157 DEGs and 3999 DEGS (with a fold change of ±1.5) in *GATA2*KD and *GATA3*KD human TSCs respectively (Fig. 6A, Supplementary Dataset S1). A comparison of DEGs revealed a striking overlap between genes that were up- or downregulated when either GATA2 or GATA3 were depleted in human TSCs (Fig. 6A). We noticed that 1910 genes were upregulated and 1517 genes were downregulated in both *GATA2*KD and *GATA3*KD human TSCs. Whereas only 362 genes were upregulated and 368 genes were downregulated specifcally in *GATA2*KD human TSCs. Additionally, 342 genes were upregulated and 230 genes were downregulated specifically in *GATA3*KD TSCs. Thus, the unbiased RNA-seq analyses indicated that GATA2 and GATA3 establish a largely similar gene expression program when promoting SynT development.

**Figure 6.**
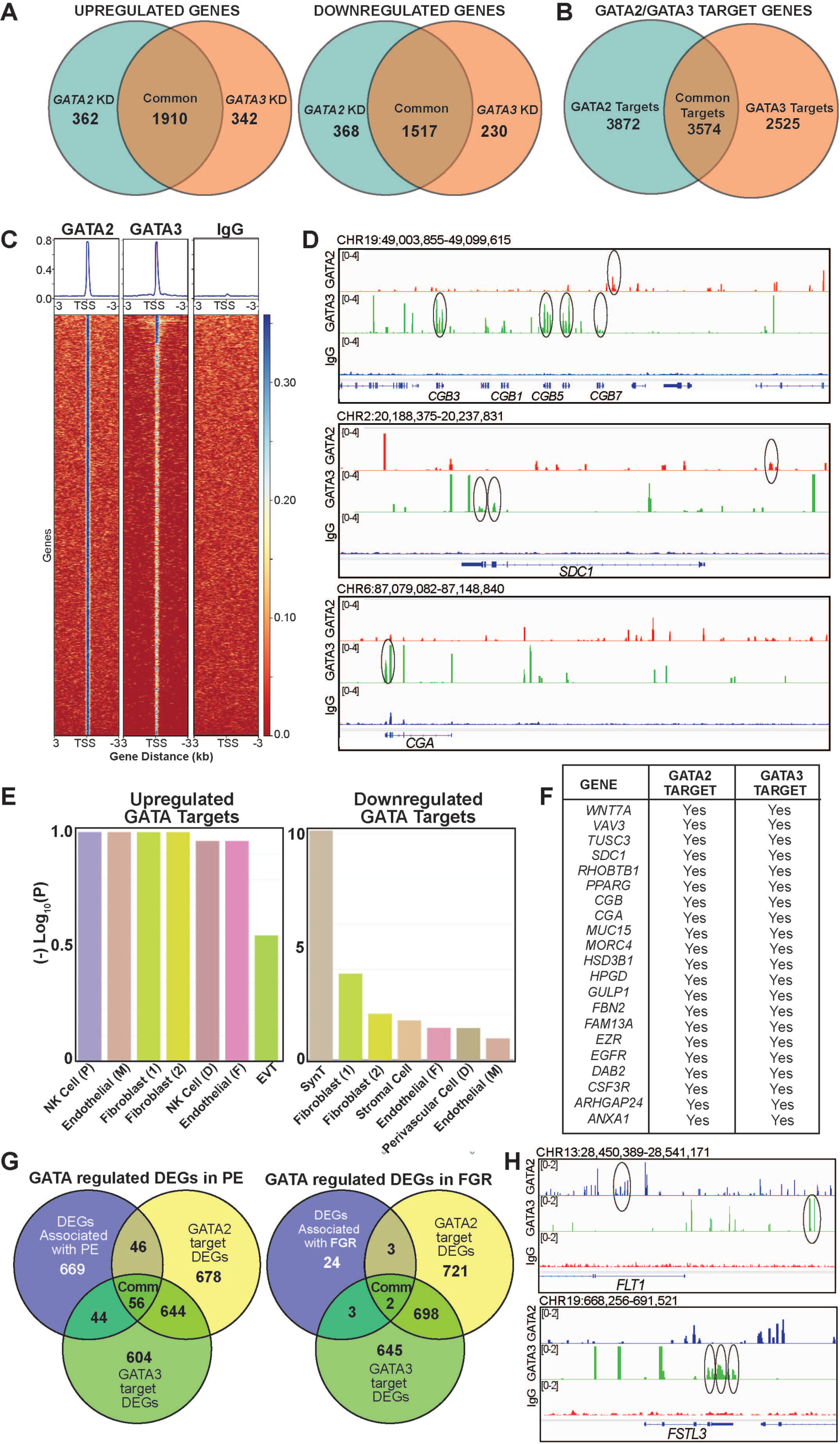
GATA2 and GATA3 directly activates SynT-specific genes. **(A)** Venn diagrams showing DEGs in GATA2KD and GATA3KD human TSCs with respect to control human TSCs when human TSCs were subjected to SynT differentiation. **(B)** Venn diagram showing number of target genes of GATA2 and GATA3 in human TSCs undergoing SynT differentiation. **(C)** Heat map representing GATA2 or GATA3 binding enrichment at the transcription start site (TSS) and 3000bp upstream and downstream of the TSS. **(D)** Integrative Genome Viewer (IGV) tracks showcasing GATA2 and GATA3 CUT&RUN Seq peaks. Statistically significant GATA2 peaks (red) and GATA3 peaks (green) are indicated at the CGB, CGA and SDC1 loci. **(E)** Placenta cell-specific gene enrichment analysis done with upregulated and downregulated genes in (A) using the PlacentaCellEnrich. Analysis highlights that the downregulated genes most signficantly represent SynT population. **(F)** Table shows a list of GATA2/GATA3 target genes, which are important for SynT differentiation and are differentially regulated in both *GATA2*KD and *GATA3*KD human TSCs. **(G)** Venn diagram showing number of genes that are significantly altered in preeclampsia (PE) and are GATA targets in human TSCs and either upregulated (26 genes) or downregulated (45 genes) when *GATA2*KD and *GATA3*KD human TSCs, when cells were subjected for SynT differentiation. Gene set curated from the placentome database (61). **(H)** IGV tracks showcasing GATA2 and GATA3 CUT&RUN Seq peaks at the *FLT1* and *FSTL3* loci.

We performed Cleavage Under Targets & Release Using Nuclease (CUT&RUN) with GATA2 and GATA3 specific antibodies to identify genes that are direct targets of GATA factors in human TSCs. Globally, we identified 8108 GATA2 binding and 5457 GATA3 binding regions (Supplementary Dataset S2). Further analyses of binding regions using Genomic Regions Enrichment of Annotations Tool (GREAT) identified 7446 putatative GATA2 traget genes and 6099 putative GATA3 target genes in human TSCs (Fig. 6B and Supplementary Dataset 2). We also noticed that 3574 genes are common targets of GATA2 and GATA3 in human TSCs (Fig. 6B). Interestingly, further analyses of GATA2 and GATA3 chromatin binding regions in common target genes revelaed a differential chromatin binding pattern for GATA2 and GATA3. The GATA2 binding regions were evenly dispersed throughout target loci (Fig. 6C). In contrast GATA3 binding regions were more concentrated near the transcription start site (TSS). Thus, our CUT&RUN analyses revealed two important aspects; (i) a large number of genes in developing SynTs are common targets of GATA2 and GATA3, and (ii) The GATA2 and GATA3 binding regions in many common target genes are different indicating that GATA2 and GATA2 might mediate unique molecular mechanisms to control expression of those genes during SynT development.

A comparative analysis between the CUT&RUN and RNA-Seq data revealed that several genes, such as *SDC1, CGA* and *CGBs,* which are specifically induced in SynTs and are differentially expressed in both *GATA2*KD and *GATA3*KD human TSCs are GATA2/GATA3 targets (Fig. 6D). We identified 355 common upregulated DEGs and 345 common downregulated DEGs, that are direct targets of both GATA2 and GATA3 (Supplementary Dataset S2). We further characterized common GATA target DEGs using placenta cell-specific gene enrichment [PlacentaCellEnrich, (60)] analysis. Interstingly the enrichment analyses revelead that the common GATA2/GATA3 target genes, which are downregulated in both *GATA2*KD and *GATA3*KD human TSCs, most significantly represent SynTs. In contrast, upregulated GATA2/GATA3 target genes most significantly represent non-trophoblast cells of a human placenta (Fig. 6E, F). Thus, our unbiased genomics analyses revealed that GATA2/GATA3-mediated gene activation, rather than gene repression is critical for human SynT development.

Pathological pregnancies, such as preeclampsia (PE) and fetal growth restriction (FGR), are often associated with altered gene expression in trophoblast cells. Therefore, we next tested association of GATA factor regulated genes with respect to pregnancies complicated by PE and FGR. We compared GATA regulated DEGs with significantly dysregulated (p≤0.05) genes in PE and FGR pregnancies identified by a recent study by Gong et al. [(61) and Supplementary Dataset S3]. We found that 56 PE-associated DEGs, including *GATA2* itself, are directly regulated by both GATA2 and GATA3 in human TSCs during SynT differentiation (Fig. 6G and supplementary Dataset S3). In addition, another 46 PE-associated DEGs are specifically regulated by GATA2 and 44 PE-associated DEGs are directly regulated by GATA3 (Fig. 6G and supplementary Dataset S3). Among the 32 significantly (p≤0.05) dysregulated FGR-associated genes [(61) and supplementary Dataset S3], two genes, Apolipoprotein B (*APOB*) and Iodothyronine Deiodinase 2 (*DIO2*), are directly regulated by both GATA2 and GATA3 during SynT differentiation (Fig. 6G and supplementary Dataset S3). In addition, three FGR-associated genes, Proteinase 3 (*PRTN3*), Prostaglandin Reductase 3 *(PTGR3), and* Triggering Receptor Expressed On Myeloid Cells 1 *(TREM1)*, are directly regulated by GATA2, whereas GATA3 directly regulates three other FGR-associated genes, *FSTL3, GPR32* and *NOG* (Fig. 6G and Supplementary Dataset S3). Intriguingly, many of the most signficantly dysregulated genes in PE [such as Fms Related Receptor Tyrosine Kinase 1 (*FLT1*), Endoglin (*ENG*), Leptin *(LEP),* and HtrA Serine Peptidase 4 *(HTRA4)*, (61, 62)] or in both PE and FGR [such as Follistatin Like 3 (*FSTL3), DIO2* and *TREM1* (61, 62)], are directly regulated in human TSCs either by GATA2 (e.g. *TREM1*) or GATA3 (e.g. *ENG, FSTL3, HTRA4* and *LEP*) or by both of them (e.g. *FLT1, DIO2*) (Fig. 6H and Supplementary Fig. S8A,B,C). Collectively, our unbiased genomics analyses in human indicated that during SynT developement GATA factors directly regulate genes, which are not only critical for SynT development, but are also implicated in pathological pregnancies, such as PE and FGR.

## Discussion

During early mouse development, GATA2 and GATA3 are specifically expressed in the trophoblast lineage and our global knockout studies revealed essential roles of GATA factors in trophoblast cells during pre-implantation and early post-implantation embryos. However, specific roles of GATA factors during SynT and labyrinth zone development has not been addressed earlier. Our in vivo studies with knockout mice along with our scRNA-seq analysis provided a better understanding of the developmental stages of SynT development from their progenitor populations and the specific role that GATA factors play in that context. We showed that there is a specific need of GATA factors to fine-tune the transcriptional program in SCA1 positive multipotent LaTP-like progenitors for their transition to committed progenitors, which will eventually develop into the terminally differentiated SynT populations to form a functional placental labyrinth (Fig. 7). Interestingly, the lack of GATA function in SCA1 positive multipotent progenitors also affected S-TGC development highlighting the important role of GATA factors during development of all of the three trophoblast subtypes of a mouse placental labyrinth. Our data also indicate that, during mouse SynT development, GATA2 and GATA3 mediate redundant roles and the function of one factor can compensate loss of the other factor so that pregnancy and embryonic develoment can progress.

**Figure 7.**
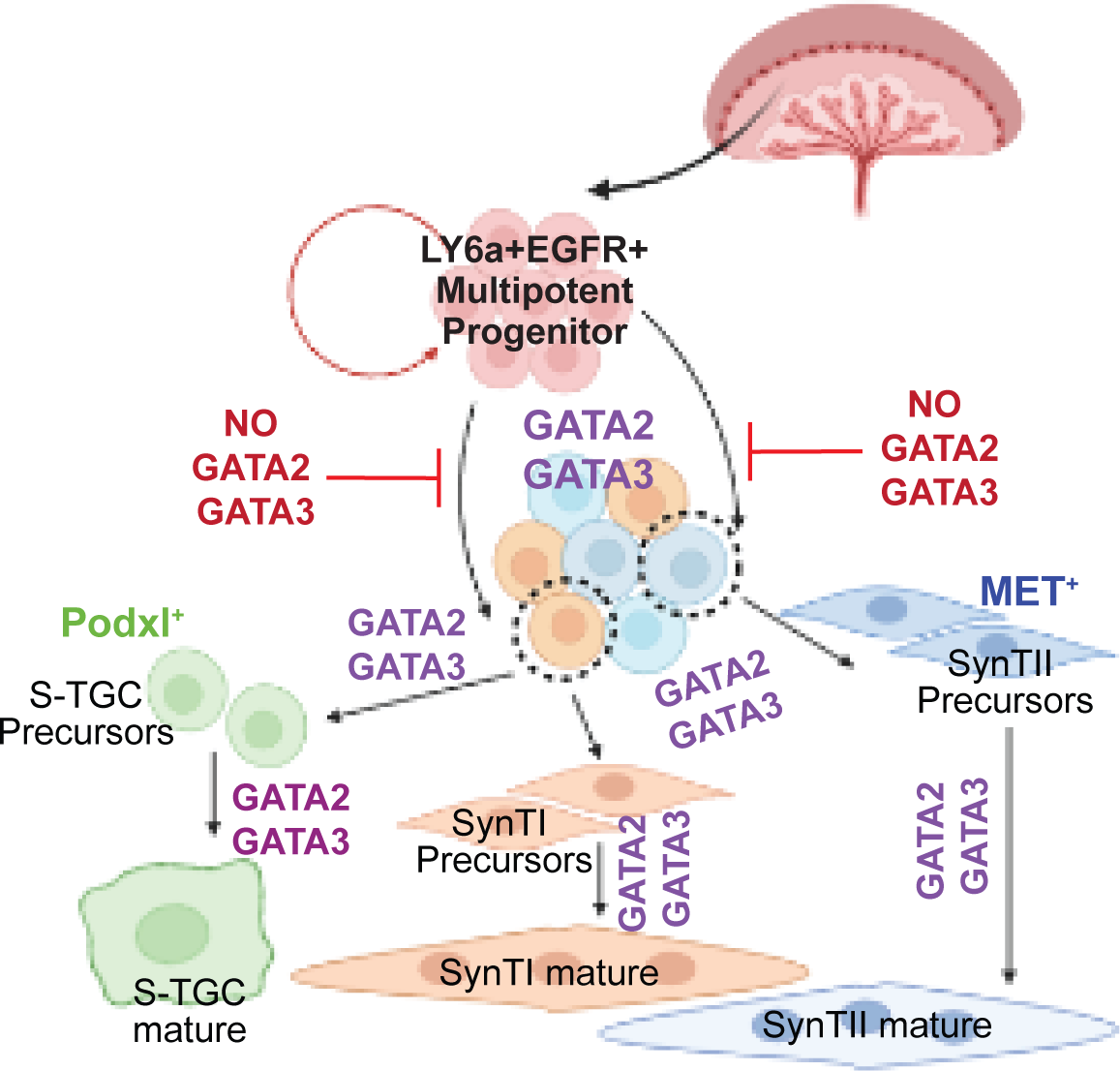
A GATA factor-dependent SynT developmental program during mouse placentation. Schematic representation of mid-gestation mouse placenta shows GATA factors regulate SynT precursor commitment and differentiation. In the presence of GATA2 and GATA3 the multipotent progenitors of the labyrinth (SCA1^+^ EGFR^+^) get committed to either the SynTI, SynTII or S-TGC lineage forming SynTI, SynTII and S-TGC precursors respectively. These precursors under the transcriptional regulation of the GATA factors differentiate into multinucleated mature SynTI and SynTII cells or S-TGCs. In the *Gcm1^Cre^*GATA-KO placentae loss of GATA factors in the labyrinth trophoblast progenitors causes then to accumulate in a multipotent progenitor state that is unable to commit to the SynT lineage.

Mouse placentae lacking three other transcription genes, *Pparg, Gcm1* and *Dlx3* as well as *c-Met* gene also showed defective SynT maturation. At the onset of labyrinth development, *Gcm1* and *Dlx3* transcriptions are induced in trophoblast progenitors. *Dlx3* is initially induced in basal EPC progenitors, which constitutes a layer in chorion and eventually differentiate to SynTI lineage (11). Later, *Dlx3* is broadly expressed in both SynTI and SynTII population. Defect in SynT development and placental labyrinth formation is also evident in the *Dlx3* knockout mice by E9.5 (22). Similarly, matured and fused SynT is absent in *Pparg* and knockout placentae (21, 63). We showed that *Dlx3* was repressed in Gcm1^Cre^ GATA DKO placentae. Furthermore, we previously showed that *Pparg and Gcm1* are also regulated by GATA factors in developing mouse placentae and *Dlx3, Pparg and Gcm1* are direct target genes of GATA2 and GATA3 in mouse TSCs (53). Thus, GATA factors seems to be the master regulator to induce *Pparg, Gcm1, Dlx3 and c-Met* during SynT maturation and labyrinth development and loss of expression of these key factors could be the molecular causes for impaired SynT commitment of multipotent labyrinth progenitors in *Gcm1^Cre^*GATA DKO placentae.

Consistent with our observation in mouse placental development, we observe a severe impairment in SynT differentiation (2D and 3D) of human TSCs due to the loss of function of either GATA2 or GATA3. This observation also indicates a difference in GATA-factor requirements for SynT differentiaton in human and mouse. Individual loss of either GATA2 and GATA3 did not affect SynT development in mouse, whereas, individual functions of either GATA2 or GATA3 seems to be essential for SynT development in human. However, unlike our studies in mouse embryos, our experiments in human TSCs does not truly represent the complexities of *in vivo* system. Thus, the *in vitro* human TSC experimental system could be more sensitive to individual loss of GATA factors.

Our RNA-seq and CUT&RUN-seq analyses revelaed that GATA2/GATA3-mediated gene activation is critical for human SynT development. Many of the SynT-specific genes are directly regulated by both of these factors and their transcription is induced by GATA factors during SynT development. However, we noticed that GATA factors are also important to mainatin self-renewal in human TSCs. Thus, it will be intersting to find out the regulatory axis that causes the switch from the self-renewal mode in the CTB progenitors towards SynT differentiation by modulating global transcriptional program via GATA factors.

In conclusion, our findings in this study establishes that GATA factors are essential for SynT development and establishment of maternal-fetal exchange interface during mammalian placentation. However, our findings in human TSCs does not indicate whether GATA factors are important for maintenance of various functions that are mediated by differentiated SynTs in a matured placenta. Also, importance of GATA factors in the context of pathological pregnancies are yet to be defined. Intriguingly, in this study we found that many of the GATA regulated genes during SynT development are dysregulated in PE and FGR. In fact, *GATA2* itself is signficantly downregulated in PE (61). Thus, it seems that there might be a direct correlation of GATA2/GATA3 transcriptional activity with dysregulated trophoblast gene expression in pathological pregnancies, such as PE and IUGR. The recent success in establishing stem cells from term placenta (64–66) and advancement of single-cell genomics analyses have provided an excellent opportunity to further study association of GATA factor transcriptional activity in the context of pathological pregnancies. We predict that the establishment of inducible genome editing system in human TSCs and studies on patient-specific stem cells from pathological pregnancies will further elucidate importance of GATA factors in the context of human pathological pregnancies.

## Materials and Methods

### Generation of conditional knockout mice strains

All procedures were performed after obtaining IACUC approvals at the Univ. of Kansas Medical Center. Male *Gata2flox/flox* (*Gata2^f/f^*)(67) mice were mated with *Tg(Gcm1-cre)1Chrn* (*Gcm1^Cre^*) female in order to generate *Gata2^f/+^; Gcm1^Cre^ ^+/wt^*. In the next step, *Gata2^f/+^; Gcm1^Cre^ ^+/wt^* female mice were bred with *Gata2^f/f^* males to generate *Gata2^f/f^; Gcm1^Cre^ ^+/wt^*. Similarly, *Gata3flox/flox* (*Gata3^f/f^*) (68) mice were used to generate *Gata3^f/f^; Gcm1^Cre^ ^+/wt^*. In the next step, *Gata2^f/f^; Gcm1^Cre^ ^+/wt^* and Gata3^f/f^; *Gcm1^Cre^ ^+/wt^* mice were crossed to generate Gata2^f/+^; *Gata3^f/+^*; *Gcm1^Cre +/wt^*. *Gata2^f/+^; Gata3^f/+^; Gcm1^Cre^ ^+/wt^* male mice were used as the *Cre* driver to restrict the expression of Cre recombinase only in the extraembryonic tissues post mating. *Gata2^f/f^; Gata3^f/f^* generated by our lab in a previous publication (69) was used as the female for the functional analysis and all future experiments.

### Immunofluorescence and Immunohistochemistry analyses

For immunostaining with mouse tissues, slides containing cryosections were dried, fixed with 4% PFA followed by permeabilization with 0.25% Triton X-100 and blocking with 10% fetal bovine serum and 0.1% Triton X-100 in PBS. Sections were incubated with primary antibodies overnight at 4°C, washed in 0.1% Triton X-100 in PBS. After incubation (1:400, one hour, room temperature) with conjugated secondary antibodies, sections were washed, mounted using an anti-fade mounting medium (Thermo Fisher Scientific, Waltham, MA) containing DAPI and visualized using Nikon Eclipse 80i fluorescent microscope. Immunohistochemistry was performed using paraffin sections of human placenta. The slides were deparaffinized by histoclear and subsequently with 100%, 90%, 80% and 70% ethanol. Antigen retrieval was done using Decloaking chamber at 80°C for 15 minutes. The slides were washed with 1X PBS and treated with 3% H2O2 to remove endogenous peroxidase followed by 3 times wash with 1X PBS. 10% goat serum was used as a blocking reagent for 1hour at RT followed by overnight incubation with 1:100 dilution of primary antibody or IgG at 4°C. Additional details are provided in SI Appendix; Supplementary Materials and Methods.

### Electron Microscopy

Placental samples were carefully isolated at E9.5. Each placenta was then placed in 2.5% glutaraldehyde fixing solution and chopped into smaller pieces to isolate regions of the labyrinth. Tissue samples were post-fixed in 1% osmium tetroxide, dehydrated in graded series of ethanol followed by dehydration with propylene oxide and infiltration with Embed 812 resin (Electron Microscopy Sciences, Hatfield, PA). Tissue polymerized in Embed 812 resin at 60°C. Blocks trimmed with EMTrim2 (Leica Biosystems, Deer Park, IL). Finally, 75nm thick sections cut with a DiATOME diamond knife using Leica UC-7 ultramicrotome and placed on 200 mesh copper grids. Sections were contrasted with uranyl acetate followed by lead citrate. Grids with sections air dried were viewed on JEOL JEM-1400 TEM at 100kV.

### Single Cell RNA-Sequencing sample preparation

Placenta samples were carefully isolated at d9.5, ensuring the decidual layer was peeled off. Individual samples were digested in the presence of collagenase (100mg/ml) at 37°C for 30 minutes, followed syringe passaging of the cells with 18G, 22G and 25G needles. 1ml of PBS+10%FBS solution is added to each tube of cell suspension to neutralize the effect of collagenase as well as to increase the volume. Finally, each placental sample was made into single-cell suspensions by passing them through a 40µm filter. Corresponding embryonic tissues were used to confirm genotypes. Based on the genotyping results the control samples were pooled together, and the knockout samples were pooled together. These samples were further processed using Debris Removal Solution and Dead Cell Removal Kit (Miltenyi Biotec., Gaithesburg, MD). Red blood cell depletion from cell suspensions were performed using anti-Mouse Ter-119 antibody (BD Biosciences, Franklin Lake, NJ). Cellular viability was checked by automated cell counter (Countess 3, Thermo Fisher Scientific) after staining cells with trypan blue. After library preparation (done by KUMC genomics core) samples were loaded into the 10X Chromium V3 platform. The raw data for scRNA-seq analyses have been submitted to the Gene Expression Omnibus (GEO) database (https://www.ncbi.nlm.nih.gov/gds), with accession No. GSE214499. Additional details of ScRNA-seq data analyses are mentioned in SI Appendix: Supplementary Materials and Methods.

### Flowcytometry analysis and cell sorting

For analyzing multipotent SCA1^+^ trophoblast population, placental single-cell suspensions were stained with PerCP/Cy5.5-conjugated anti-mouse CD34 (BioLegend, San Diego, CA), PE-conjugated anti-mouse Ly-6A/E (SCA1) (BioLegend), PE/Cy7 anti-mouse CD45 (BioLegend) and Pacific, Blue-conjugated anti-mouse Lineage cocktail (BioLegend) monoclonal antibodies. Unstained, isotype and single-color controls were used for optimal gating strategy. Samples were run on either an LSRII flow cytometer or an LSRFortessa (BD Biosciences), and the data were analyzed using FACSDiva software.

### Collection and analysis of human placentae

First trimester, pre-term and term placental samples from normal and pathological pregnancies were obtained from RCWIH biobank, Toronto, or collected at the University of Kansas Medical Center after IRB approval from both the universities. The tissues we embedded in OCT for cryosectioning or used for RNA isolation.

### Human TSC culture

Human TSC lines were derived from first trimester CTBs, as described earlier (58). To maintain stem state culture, human TSCs were cultured on collagen IV-coated (5µg/ml) plate in DMEM/F12 medium, supplemented with 0.1 mM 2-mercaptoethanol, 0.2% FBS, 0.5% Penicillin-Streptomycin, 0.3% BSA, 1% ITS-X supplement, 1.5 µg/ml L-ascorbic acid, 50 ng/ml EGF, 2 µM CHIR99021, 0.5 µM A83-01, 1 µM SB431542, 0.8 mM Valproic acid and 5 µM Y27632. For SynT(2D) differentiation, plates were coated with 2.5 µg/ml of collagen IV, TSCs were then cultured in DMEM/F12 medium, supplemented with 0.1 mM 2-mercaptoethanol, 0.5% Penicillin-Streptomycin, 0.3% BSA, 1% ITS-X supplement, 2.5 mM Y27632, 2 mM forskolin, and 4% KSR. For, SynT(3D) differentiation, no plate coating was required. Cells were cultured in the same media used for SynT(2D) with 50ng/ml of EGF.

### Short hairpin RNA (shRNA) mediated RNAi

To generate *GATA2* or *GATA3* knockdown human TSCs, lentivirus particles carrying shRNA against *GATA2* mRNA with sequence (GTGCAAATTGTCAGACGACAA) or *GATA3* mRNA with sequence (TCTGGAGGAGGAATGCCAATG) were used. A scramble shRNA with sequence (CCTAAGGTTAAGTCGCCCTCGC) was used as control. Transduced cells were selected in the presence of puromycin (1.5-2µg/ml). Selected cells were tested for knockdown efficiency and used for further experimental analyses.

### RNA-Seq analyses

RNA-seq analysis was performed according to published protocol(69, 70). Total RNA from the control human TSCs as well as GATA2-KD and GATA3-KD human TSCs were isolated using RNeasy Mini Kit (Qiagen, Germantown, MD) per the manufacturer’s protocol with on-column deoxyribonuclease digestion. RNA concentrations were quantified using a NanoDrop Spectrophotometer at a wavelength of 260 nm. Integrity of the total RNA samples was evaluated using an Agilent Technologies 2100 Bioanalyzer. The total RNA fraction was processed by oligo dT bead capture of mRNA, fragmentation, and reverse transcription into cDNA. After ligation with the appropriate Unique Dual Index (UDI) adaptors, the cDNA library was prepared using the Universal Plus mRNA-seq +UDI library preparation kit (NuGEN; Marion, SD). The raw data for RNA-seq analyses have been submitted to the Gene Expression Omnibus (GEO) database (https://www.ncbi.nlm.nih.gov/gds), with accession No.GSE214486.

### Statistical analyses

Statistical significance was determined for quantitative RT-PCR analyses for mRNA expression and for FACS analyses. We performed at least n = 3 experimental replicates for all these experiments. For statistical significance of generated data, statistical comparisons between two means were determined with Student’s t test, and significantly altered values (P ≤ 0.01) are highlighted in the figures by an asterisk. RNA-seq data were generated with n = 3 experimental replicates per group. The statistical significance of altered gene expression (absolute fold change ≥2.0 and false discovery rate (FDR) q-value ≤0.05) was initially confirmed with right-tailed Fisher’s exact test. Independent data sets were analyzed using GraphPad Prism software.

#### GEO database accession codes for sequencing data

The raw genomics data are submitted to GEO data base.Data for Single-cell RNA sequencing of E9.5 control and *Gcm1^Cre^*GATA DKO mouse placentae is submitted to with GEO accession number GSE214389. Raw data for bulk RNA-seq and CUT & RUN seq in human TSCs are submitted with GEO accession numbers, GSE214634 and GSE214486, respectively.

**Additional Materials and Methods including details of organoid cultures and CUT & RUN analyses are mentioned in SI Appendix: Supplementary Materials and Methods.**

## ACKNOWLEDGEMENTS

This research was supported by NIH grants HD062546, HD103161, HD102188 and HD101319 to Soumen Paul. This study was also supported by administrative support from the IRDS, KUMC and various core facilities, including the Genomics Core, Transgenic and Gene Targeting Institutional Facility, Imaging and Histology Core and the Flow Cytometry Core. We thank Drs. Hiroaki Okae and Takahiro Arima of Tohoku University Graduate School of Medicine, Japan, for sharing human TSC lines. We thank Dr. Pratik Home for his intellectual support about the research project and Ms. Brandi Miller for critical comments on the manuscript.

