## Supplemental text and Figures for "GATA-regulated transcriptional program dictate cell fate equilibrium to establish the maternal-fetal exchange interface and fetal development"

**The PDF file includes:**
(i) Supporting Materials and Methods

(ii) Supplementary Figures, Fig. S1- S8
(iii) SI References

**Other supporting materials for this manuscript include the following:**

Datasets S1 to S3

#### Supporting Materials and Methods

##### Collection of mouse embryos and tissue isolation

Animals were euthanized on at desired day points, as indicated in the main text. Pregnant female animals were identified by presence of vaginal plug (gestational day 0.5), and embryos were harvested at various gestational days. Uterine horns from pregnant females were dissected out, and individual embryos were analyzed under microscope and photographed. Conceptuses were dissected to isolate embryos, yolk sacs, and placentae. All embryos and placentae were photographed at equal magnification for comparison purposes. Uteri containing placentation sites were dissected from pregnant female mice on E8.5-E14.5., tissues for histological analysis were kept in dry ice cooled heptane and stored at −80 °C. Tissues were subsequently embedded in optimum cutting temperature (OCT) (Tissue-Tek, Torrance, CA) and were cryosectioned (10mm thick) for immunohistochemistry (IHC) studies using Leica CM-3050-S cryostat. Yolk sacs from each of the dissected embryos were collected, and genomic DNA preparation was done using Extract-N-Amp tissue PCR kit (Sigma, St. Louis, MO). Placenta tissues were collected in RLT buffer, and RNA was extracted using RNAeasy Mini Kit (Qiagen). RNA was eluted and concentration was estimated using Nanodrop ND1000 spectrophotometer. Placenta samples were carefully isolated, ensuring the decidual layer was peeled off. Individual samples were briefly digested in the presence of collagenase and were made into single-cell suspensions by passing them through a 40μm filter. These cell suspensions were further used for Flow analysis or FACS. Corresponding embryonic tissues were used to confirm genotypes.

##### Genotyping

Genomic DNA samples were prepared using tail tissues or embryonic tissues from the mice using the REDExtract-N-Amp Tissue PCR kit (Sigma-Aldrich). Genotyping was done using REDExtract-N-Amp PCR ReadyMix (Sigma-Aldrich) and respective primers. Respective primers are listed in the materials and methods section.

##### Immunofluorescence and Immunohistochemistry analyses

For immunostaining with mouse tissues, slides containing cryosections were dried, fixed with 4% PFA followed by permeabilization with 0.25% Triton X-100 and blocking with

10% fetal bovine serum and 0.1% Triton X-100 in PBS. Sections were incubated with primary antibodies overnight at 4°C, washed in 0.1% Triton X-100 in PBS. After incubation (1:400, one hour, room temperature) with conjugated secondary antibodies, sections were washed, mounted using an anti-fade mounting medium (Thermo Fisher Scientific) containing DAPI and visualized using Nikon Eclipse 80i fluorescent microscope. *mT/mG* positive embryos and cryosections were imaged directly under a Nikon Eclipse 80i fluorescent microscope. Immunohistochemistry was performed using paraffin sections of human placenta. The slides were deparaffinized by histoclear and subsequently with 100%, 90%, 80% and 70% ethanol. Antigen retrieval was done using Decloaking chamber at 80$^{O}$C for 15minutes. The slides were washed with 1X PBS and treated with 3% H2O2 to remove endogenous peroxidase followed by 3 times wash with 1X PBS. 10% goat serum was used as a blocking reagent for 1hour at RT followed by overnight incubation with 1:100 dilution of primary antibody or IgG at 4$^{O}$C. The slides were washed with 1X PBS and 1:200 dilution of secondary antibody was used for 1 hour at RT. The slides were washed again with 1X PBS followed by treatment with horseradish peroxidase streptavidin for 20 minutes at RT. The slides were washed again and proceeded to color development using DACO 1ml buffer and 1 drop of chromogen. The reaction was stopped in distilled water after sufficient color developed. The slides were counterstained with Mayer's hematoxylin for 5minutes and washed with warm tap water until sufficient bluish coloration observed. The slides were then dehydrated by sequential treatment using 70%, 80%, 90%, 100% ethanol and histoclear. The sections were completely dried and mounted using Toluene as mounting agent and imaged using Nikon TE2000 microscope.

**Single-cell RNA sequencing data analysis**

The sequencing results were loaded using Read10X function of Seurat and processed through Cellranger pipeline from 10X genomics to obtain unique molecular identified (UMI) count matrix. The values in this matrix represent the number of molecules for each feature (i.e., gene; row) that are detected in each cell (column). In the next step we have created a Seurat object for individual samples using the CreateSeuratObject function and provided parameters to perform some initial filtering in order to exclude genes that are expressed in fewer than 3 cells, and to exclude cells that contain fewer than 500 expressed genes. Next, we start with filtration of cells based on QC metrics, which is followed by data normalization, scaling and finally detection of highly variable features.

We excluded the cells having less than 500 and greater than 5000 unique genes. Along with it, we also excluded nuclei having mitochondrial counts 10 percentage for further downstream analysis. We put all samples into list and used NormalizeData() and ScaleData() function of Seurat with default settings for normalizing each of sample. Further we used FindVariableFeatures function and provided 2500 features to subset the samples for downstream analysis. SelectIntegrationFeatures function screens the features that will be used while performing integration and FindIntegrationAnchors was implemented to get anchors. Finally IntegrateData function was used which create integrated assay. We used DefaultAssay to make this integrated assay default, which will be used in downstream analysis. We used RunPCA function to perform PCA. Next characterization of the principal components and estimate the number of significant PCs that captures the signal of interest, while minimizing noise was accessed using ElbowPlot and DimHeatmap functions. Further functions like FindNeighbors and FindClusters were executed sequentially for clustering. In the final steps dimensional reduction methods UMAP and tSNE was performed using RunUMAP and RunTSNE function of Seurat. Differentially expressed genes were identified using FindAllMarkers and cell clusters were identified based on the expression of marker genes mentioned in Supplementary Table 1 below. To further analyze trophoblast cell cluster, data from initial clustering was further analyzed for trophoblast cell clusters using the function SubsetData based upon annotations from marker genes identified by FindAllMarkers. Integration of the trophoblast only dataset was performed using the same integration methods above (using 20 dimensions for FindIntegrationAnchors and IntegrateData, and 20 PCs and a resolution of 0.5 for FindClusters and RunUMAP). Differentially expressed genes for each integrated dataset were identified using FindAllMarkers (Supplementary Table 2).

**Supplementary Table 1: Marker Genes for Initial Cell Clustering**

| | | |
|---|---|---|
| Trophoblast | 2,4,13,14 | Epcam, Ly6a, Prl3d1, Prl2c2, Tpbpa |
| Endothelial | 3 | Pecam1, Cldn5, Cd34, Kdr, Eng |

| | | |
|---|---|---|
| Endodermal | 6, 19 | Procr, Cldn6, Rbp4, Afp Procr, S100g, Klf5 |
| Blood cells | 0,15,17- B-cell | Cd74, Igkc, Cd79a, Cd79b |
| | 9 - T-cell | Trbc1, Lmo2 |
| | 8 - NK cells | Thy1, Klrg1, Cst7, Prf1 |
| | 1,20 - Erythroblasts | Blvrb, Hbb-y |
| | 11 - Megakaryocytes | Itga2b, Tal1, Nfe2, Lmo2 |
| | 7 - Macrophages | Spp1, Adgre1, Cx3cr1, Apoe |
| | 21 - Dendritic cells | Cd83, Cst3, H2-Aa, Cd209a |
| | 12 - granulocytes | Neat1, Cd69 |
| Stromal | 18,10,16,5 | Acta2, Dcn, Col3a1, Col1a2, Adm, Cryab |

**Supplementary Table 2: Marker Genes for Trophoblast Cell Clusterinng**

| **Trophoblast sub-clusters** | **Cluster number** | **Marker genes** |
|---|---|---|
| Multipotent progenitors | 3 | *Tcf7l1, mKi67, Tbx3, Ly6a, Egfr* |
| SynTI precursors | 6 | *Slc16a1, Slc40a1, Nr6a1, Dlx3* |
| SynTII precursors | 5 | *Slc16a3, Esx1, Fabp3, Wnt7b* |
| S-TGC precursors | 8 | *Podxl, Lifr, Hsd17b2* |
| JZ precursors 1 (JZP1) | 1 | *Hand1, Ctsd, Podxl, Prl2b1* |
| JZ precursors 2 (JZP2) | 2 | *Prl8a1, Slco2a1* |
| JZ precursors 3 (JZP3) | 7 | *Prune2, Prl2b1, Prl3b1, Prl8a9* |
| JZ precursors 4 (JZP4) | 4 | *Pla2g4d, Ncam1, Igfbp7, Pcdh12, Tpbpa* |

**RNA velocity study**

RNA Velocity enables measuring the underlying kinetics of a gene's expression from each cell based on the single cell RNA sequencing dataset. The concept of RNA velocity utilizes the information about newly transcribed pre-mRNAs (unspliced) from mature mRNAs (spliced), which can be detected in standard single-cell RNA-seq protocols from the presence of introns. The RNA velocity function distinguishes the spliced from the unspliced mRNA for each gene in the cell and generates velocities across each gene based on their expression. The change in mRNA abundance is termed RNA velocity (1). Positive velocity indicates a recent increase in pre-mRNA transcripts (thus abundances being higher than expected in steady state) followed by up-regulation in spliced transcripts. Conversely, negative velocity indicates down-regulation. The combination of velocities across genes is then used to estimate the future state of an individual cell. For the study of RNA velocity in single cell transcriptomes data, we generated the loom files which contains both splices and un-spliced contents using the Space Ranger output along with the GTF file. The tool velocyto 0.17.16 has been utilized to generate loom file. Further, we have used scVelo tool to perform RNA velocity on our sample. The loading was done using scv.read function. In the next step, we perform the gene selection, normalizing and logarithmizing using scv.pp.filter_and_normalize function. In the next step Scanpy 1.9.1 was used to get the first and second order moments (means and uncentered variances) calculated using scv.pp.moments, which are required for velocity estimation and stochastic estimation, respectively. We have used stochastic model of scVelo to calculate the RNA velocity for both samples. The tSNE cell embeddings were utilized to display the RNA velocity.

**Single Cell Proportion test**

The variability between biological samples used for scRNA-Seq can be high due to variability in the source of the samples, technical reasons like the dissociation protocol used or intrinsic properties of the sample, the single cell proportion test is a function of the R library (https://github.com/rpolicastro/scProportionTest) that analyses scRNA-Seq samples to account for variability in the proportions of cell clusters between samples. A permutation test first calculates a p-value for each cluster, then bootstrapping is used to calculate the confidence interval for the magnitude difference.

**Quantitative RT-PCR**

Total RNA from either whole cell extract or placental tissues was isolated using RNeasy Mini Kit (Qiagen, 74104) according to manufacturer protocol. Purified RNA was used to prepare cDNA using a cDNA preparation kit and analyzed by qRT-PCR following procedures described earlier (2)_ENREF_66. The mouse or human specific primers used in qRT-PCR are listed below. 20ng equivalent of cDNA was used for amplification reaction using Power SYBR Green PCR master mix (Applied Biosystems, Waltham, MA).

**Western Blot analysis**

Cell pellets were washed once with PBS followed by addition of 1X SDS-PAGE buffer for protein extract preparation and western blot analyses were performed following earlier described protocol(2). Antibodies used for the study are mentioned below.

**Organoid culture:**

Human trophoblast cell (human TSC) lines that were derived from first trimester CTBs were used to generate the organoids. We followed earlier described protocols that were used to generate CTB-organoids (5-7). Human TSCs were cultured and passaged in DMEM/F12 media that was supplemented with 0.1mM 2-mercaptoethanol, 0.2% FBS, 0.5% penicillin-streptomycin, 0.3% BSA, 1% ITS-X supplement, 1.5μg/ml L-ascorbic acid, 50ng/ml EGF, 2μMCHIR99021, 0.5μM A83-01, 1μM SB431542, 0.8mM VPA and 5μM Y27632. Cells were cultured at 37°C in 5% CO2. To generate GATA2 KD and GATA3 human TSCs, shRNA mediated RNAi was performed as described earlier. To obtain organoids from human TSCs (control/ GATA2 KD human TSCs/ GATA3 KD human TSCs) we modified the protocol described previously (7). The hTS cells were harvested and re-suspended in ice-cold basic trophoblast organoid medium (b-TOM) containing advanced DMEM/F12 supplemented with 10mM HEPES, B27 (Gibco, Waltham, MA), N2 (Gibco) and 2mM glutamine (Gibco). The cells were then centrifuged for 3 minutes at 1200rpm following which the cells were re-suspended in ice-cold advanced trophoblast organoid medium (aTOM) which is bTOM supplemented with 100ng/ml R-spondin (PeproTech, Rocky Hill, NJ), 1μM A83-01 (Sigma), 100ng/ml recombinant human epidermal growth factor (rhEGF, Wako), 50ng/ml recombinant murine hepatocyte growth factor (rmHGF, PeproTech), 2.5μM prostaglandin E2 (R&D System, Minneapolis, MN), 3μM CHIR99021 (Wako, Richmond, VA) and 100ng/ml

Noggin (Invitrogen, Waltham, MA). Growth factor reduced Matrigel (Corning) was added to the a-TOM cell suspension to reach a final concentration of 60%. 35μl of the viscous cell solution containing 2.5x104 cells was plated in the center of a 24-well plate. The solution rests as a dome-shaped droplet in the center of the well. The plates are then turned upside down and kept at 37°C for 10-15 minutes to ensure proper spreading of the cells in the solidifying matrigel domes. Finally, the plates are returned to their upward position and the domes are overlaid with 500μl of room temperature a-TOM medium. The organoids are allowed to form for 10 days with fresh media being changed every 2 days. Brightfield images were taken to observe the growth of the organoids. For staining purposes, the organoid growth was stopped at day 10 by removing the a-tom media and fixing them in ice cold 4% paraformaldehyde solution. The fixed organoids were then embedded in 4-5% agarose followed by making a paraffin block. These paraffinized blocks were then sectioned into 10μm sections using a microtome and stained (staining protocol and mentioned above).

**CUT & RUN Analyses**

Cells were harvested from culture, 200,000 live cells were taken per sample (control HTS cells) and spiked with 20% spike-in cells (freshly harvested HEK293 cells). The spiked cell suspensions were then washed twice with the wash buffer (1 ml 1M HEPES pH 7.5, 1.5ml 5M NaCl, 12.5μL 2M Spermidine, bring the final volume to 50ml with $dH_2O$, and add 1 Roche Complete Protease Inhibitor EDTA-Free tablet). After washing, 10μl of Concanavalin-A coated beads were added to 1ml of each sample and incubated under rotation at room temperature for 10minutes. Using a magnetic stand the cells were separated from the suspension, the liquid was discarded, and the cells were resuspended in 100μl of antibody binding buffer (400μL 1M HEPES-KOH pH 7.9, 200μL 1M KCl, 20μL 1M $CaCl_2$ and 20μL 1M $MnCl_2$, and bring the final volume to 20 ml with $dH_2O$). Primary antibody of interest (GATA2 or GATA3 or IgG antibody; 1:100 dilution) was added to the samples and incubated overnight at 4°C with intermittent shaking. Tri-Methyl-Histone H3 (Lys4) (C42D8) was used as a positive control antibody. The samples were washed twice with 150μL of Dig-wash buffer (200μL 5% Digitonin with 20 ml Wash buffer) while keeping the samples on ice, magnetic stand was used to remove the residual liquid. Next, A/G-MNase enzyme (~700 ng/ml diluted Dig-wash buffer) was added to each sample and incubated for 1 hour with intermittent shaking/rotation in cold room. Incubation was followed by vertexing, a quick spin and then placed on the magnet stand to separate out and discard the liquid. The samples were washed twice with Dig-wash buffer. For the next few steps an ice-water bath is made and a metal eppendorf rack is placed in the ice-cold water bath. 100mM $CaCl_2$ (Thermo Fisher Scientific) solution, Dig-wash buffer and all the samples were placed on the metal blocks kept in ice water. 2µL 100 mM $CaCl_2$ was then added to each sample, mixed well and the tubes were immediately replaced in the 0°C block for the digestion time (30 min). 2X STOP buffer (3.44ml $dH_2O$ add 136µl 5M NaCl, 400µL 0.2M EGTA, 40µL 5% Digitonin, 10µL RNase A, 20µL 20 mg/ml glycogen) is prepared during this incubation period. 100µL 2X STOP buffer is then added, mixed by gentle vortexing and incubated for 10 min 37°C to release CUT&RUN fragments from the insoluble nuclear chromatin. Samples are then centrifuged for 5 min in 4°C at 16,000 x g and placed on magnetic stand. Transfer the supernatant to a fresh 1.5-ml microcentrifuge tube. 2µL of 10% SDS (to 0.1%) and 2.5µL of Proteinase K (Thermo Fisher Scientific) (20 mg/ml)/sample was added, mixed by inversion, and incubate 10 min 70°C. 300µL of Phenol–chloroform–isoamyl alcohol 25:24:1 (Invitrogen) was added and mixed by vortexing at full speed. The solution was transferred to a phase-lock tube (Qiagen), centrifuged for 5 min in room temperature at 16,000 x g in fixed angle centrifuge. Next, 300 µL chloroform was added and inverted ~10x times to mix, then centrifuged for 5 min at room temperature at 16,000 x g in fixed angle centrifuge. Removed liquid by pipetting to a fresh tube. 750µL of 100% ethanol was added and mixed by tube inversion, chilled on ice for 10 min and centrifuged for 10 min at 4 °C 16,000 x g in fixed angle centrifuge. Excess liquid was poured off and drained on a paper towel and left to air dry. When the pellet is dry, it was dissolved in 20µL 1 mM Tris-HCl pH8 0.1 mM EDTA. The DNA was quantified using 1µL of the freshly dissolved pellet, for example using fluorescence detection with a Qubit instrument (Life Technologies, Waltham, MA). Evaluated the presence of cleaved fragments and the size distribution by capillary electrophoresis with fluorescence detection using a Tapestation instrument. For Library preparation Swift bioscience 1S Plus Combinatorial Dual Indexing Kit and Accel-NGS 1S Plus DNA Library Kit protocol was followed. We have used “nf-core” pipeline (3) for downstream analyses of CUT & RUN data.

**Heatmap for binding enrichment**

We have used deepTools (4) to see the binding enrichment of GATA2 or GATA3 around the transcription start site (TSS). For this we first calculate the values for heatmaps by providing the parameters "-a 3000, -b 3000 --missingDataAsZero -p 20 –skipZeros" to the function "computeMatrix reference-point" of deepTools, along with the bigWig file and bed file containing peaks coordinates. The output of this matrix was supplied to the plotHeatmap function with parameter "--sortRegions descend" to generating the heatmap around TSS.

**Primers used for genotyping:**

| | | |
|---|---|---|
| Gata2$^{f/f}$ and *Gata2-KO* | GCCTGCGTCCTCCAACACCTCTAA | TCCGTGGGACCTGTTTCCTTAC |
| *Gata3$^{f/f}$* | CAGTCTCTGGTATTGATCTGCTTCTT | GTGCAGCAGAGCAGGAAACTCTCAC |
| *Gata3-KO* | TCAGGGCACTAAGGGTTGTTAACTT | GTGCAGCAGAGCAGGAAACTCTCAC |
| *Cre* | AAAATTTGCCTGCATTACCG | ATTCTCCCACCGTCAGTACG |

#### List of RT-PCR primers

| Human gene | Forward | Reverse |
|---|---|---|
| *GATA2* | *CCAGCTTCACCCCTAAGCAG* | *CCACAGTTGACACACTCCCG* |
| *ERVW-1* | *CTACCCCAACTGCGGTTAAA* | *GGTTCCTTTGGCAGTATCCA* |
| *CGA* | *TCTGGTCACATTGTCGGTGT* | *TTCCTGTAGCGTGCATTCTG* |
| *CGB* | *GTGTGCATCACCGTCAACAC* | *GGTAGTTGCACACCACCTGA* |
| *PSG4* | *CGATGGGACTGGAGGAGTAA* | *AGTTGCTGCTGGAGATGGAG* |
| *TEAD4* | *ACG GCC TTC CAC AGT AGC AT* | *CTT GCC AAA ACC CTG AGA CT* |
| *VGLL1* | *TCA GAG TGA AGG TGT GAT GCT* | *GCA CGG TTT GTG ACA GGT ACT* |
| *TP63* | *GTC ATT TGA TTC GAG TAG AGG GG* | *CTG GGG TGG CTC ATA AGG T* |
| *PPARG* | *ACC AAA GTG CAA TCA AAG TGG A* | *ATG AGG GAG TTG GAA GGC TCT* |

| | | |
|---|---|---|
| *ITGA6* | *GGC GGT GTT ATG TCC TGA GTC* | *AAT CGC CCA TCA CAA AAG CTC* |
| *TFAP2C* | *TAC TGG GAG GTG TTC TCA GAA G* | *GGC CGG AAG ATT CAA CCC AAT* |
| *HPRT1* | *ACCCTTTCCAAATCCTCAGC* | *GTTATGGCGACCCGCAG* |
| *18SrRNA* | *AACCCGTTGAACCCCATT* | *CCATCCAATCGGTAGTAGCG* |
| **Mouse gene** | **Forward** | **Reverse** |
| *Gcm1* | *AGAGATACTGAGCTGGGACATT* | *CTGTCGTCCGAGCTGTAGATG* |
| *Dlx3* | *CACTGACCTGGGCTATTACAGC* | *GAGATTGAACTGGTGGTGGTAG* |
| *Pparg* | *AGCTGTCATTATTCTCAGTGGAG* | *ATGTCCTCGATGGGCTTCAC* |
| *SynA* | *ATGGTTCGTCCTTGGGTTTTC* | *GTGTTGAGTGAGGTTTACCAGG* |
| *SynB* | *TGGGTCCTCTGTTTCGTCCTT* | *GGGAAGAGTTGGTATCACGTAGG* |
| *Ctsq* | *CATTGCCAGTTGACAACACCAG* | *ATAGCCTTCATTTCGCCAATCA* |
| *Hand1* | *CTACCAGTTACATCGCCTACTTG* | *ACCACCATCCGTCTTTTTGAG* |
| *Prl2c2* | *TCCTGGATACTGCTCCTACTACT* | *GACCATTCCTCATTGCACACA* |
| *Tpbpa* | *TCCGGTCAGCTAACTGATGA* | *TCCTCTTCAAACATTGGGTGT* |
| *Prl3d1* | *ACATTTATCTTGGCCGCAGATGTGT* | *TTTAGTTTCGTGGACTTCCTCTCGAT* |
| *18SrRNA* | *AGTTCCAGCACATTTTGCGAG* | *TCATCCTCCGTGAGTTCTCCA* |

**Antibodies used for Immunofluorescence, Immunohistochemistry and Western Blots**

| Primary Antibodies | company | Catalog number |
|---|---|---|
| GATA2 | Abcam | ab109241 |
| GATA3 | BD Biosciences | 558686 |
| Pan-cytokeratin | Abcam | ab9377 |
| β-actin | Sigma | A5441 |

| | | |
|---|---|---|
| HCGβ | Abcam | ab53087 |
| Cytokeratin 7 | Dako | M7018 |
| E-cadherin | Abcam | ab1416 |
| MCT1 | EMD Millipore | AB1286-I |
| MCT4 | EMD Millipore | AB3314P |
| Proliferin | Santa Cruz | sc-47347 |
| **Secondary Antibodies** | | |
| Alexa fluor 488 goat anti-rabbit IgG | Invitrogen | A11008 |
| Alexa fluor 568 goat anti-mouse IgG | Invitrogen | A11031 |
| Alexa fluor 488 donkey anti-mouse IgG | Invitrogen | A21202 |
| Alexa fluor 568 donkey anti-rabbit IgG | Invitrogen | A10042 |
| Alexa fluor 568 donkey anti-goat IgG | Invitrogen | A11057 |
| Alexa fluor 488 goat anti-chicken IgY | Invitrogen | A32931 |
| Goat anti-mouse IgG-HRP | Santa Cruz | sc2005 |
| Goat anti-rabbit IgG-HRP | Santa Cruz | sc2004 |

| Antibodies used in FACS | Species raised in | Vendor | Catalog number |
|---|---|---|---|
| APC anti-mouse CD117 (c-kit) | Rat | BioLegend | 105812 |
| PerCP/Cy5.5 anti-mouse CD34 | American Hamster | BioLegend | 128607 |
| PE anti-mouse Ly-6A/E (Sca-1) | Rat | BioLegend | 108108 |
| PE/Cy7 anti-mouse CD45 | Rat | BioLegend | 103113 |
| Pacific Blue anti-mouse Lineage Cocktail | Rat | BioLegend | 133310 |

**Fig. S1**

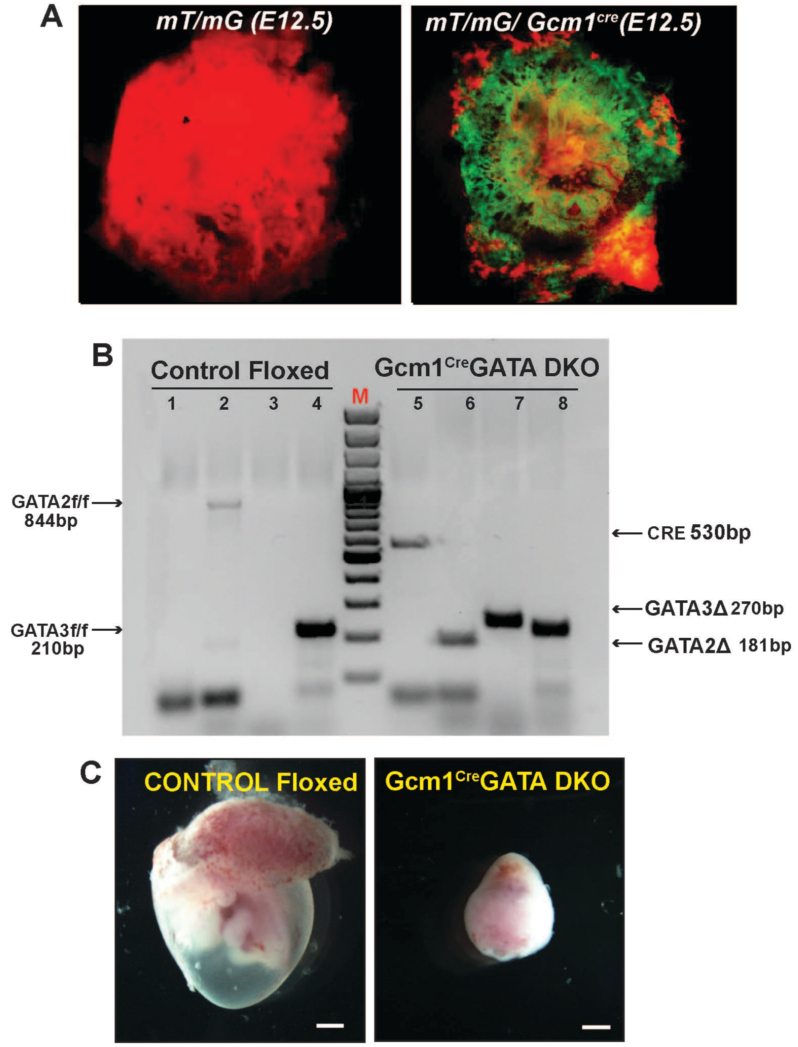

**D**

| MOUSE PHENOTYPE ANALYSIS | | | | |
|---|---|---|---|---|
| | Conceptuse analyzed | | Embryos with Developmental Defect | |
| | Floxed Control | Gcm1Cre GATA DKO | Control | Gcm1Cre GATA DKO |
| E8.5 | 18 | 10 | 0 | 0 |
| E9.5 | 40 | 28 | 0 | 27 |
| E10.5 | 27 | 13 | 0 | 13 |

**Figure S1.** (A) (left) Only mT/mG placenta at E12.5 fluoresces red. (Right) mT/mG/ *Gcm1* $^{Cre}$ placenta at E12.5, shows the specific expression of the Cre recombinase (green fluorescence) persists only in the developing labyrinth. (B)Agarose gel image representing the genotyping analysis. Homozygous *Gcm1* $^{Cre}$GATA DKO (*Gata2* $^{f/f}$ *Gata3* $^{f/f}$ *Gcm1* $^{Cre}$) embryos were confirmed by genotyping. The control samples have no Cre band (lane1), no GATA3 deletion band (lane 3), show floxed band for GATA2 (lane 2) and GATA3(lane4). The *Gcm1* $^{Cre}$GATA DKO samples show prominent Cre band (lane 5), deletion band for GATA2 (lane 6) and GATA3 (lane 7) and GATA3 floxed band (lane 8). M, molecular weight marker. (C) Images show E10.5 control and *Gcm1* $^{Cre}$GATA DKO conceptuses. Most of the *Gcm1* $^{Cre}$GATA DKO embryos are resorbed at E10.5 (scale bars 500 μm) . (D) Table showing number and phenotype of control and *Gcm1* $^{Cre}$GATA DKO embryos, analyzed at E8.5 (n=28), E9.5 (n=68) and E10.5 (n=40), 'n' represents number of independent experiments conducted with multiple pregnant females.

**Fig. S2**

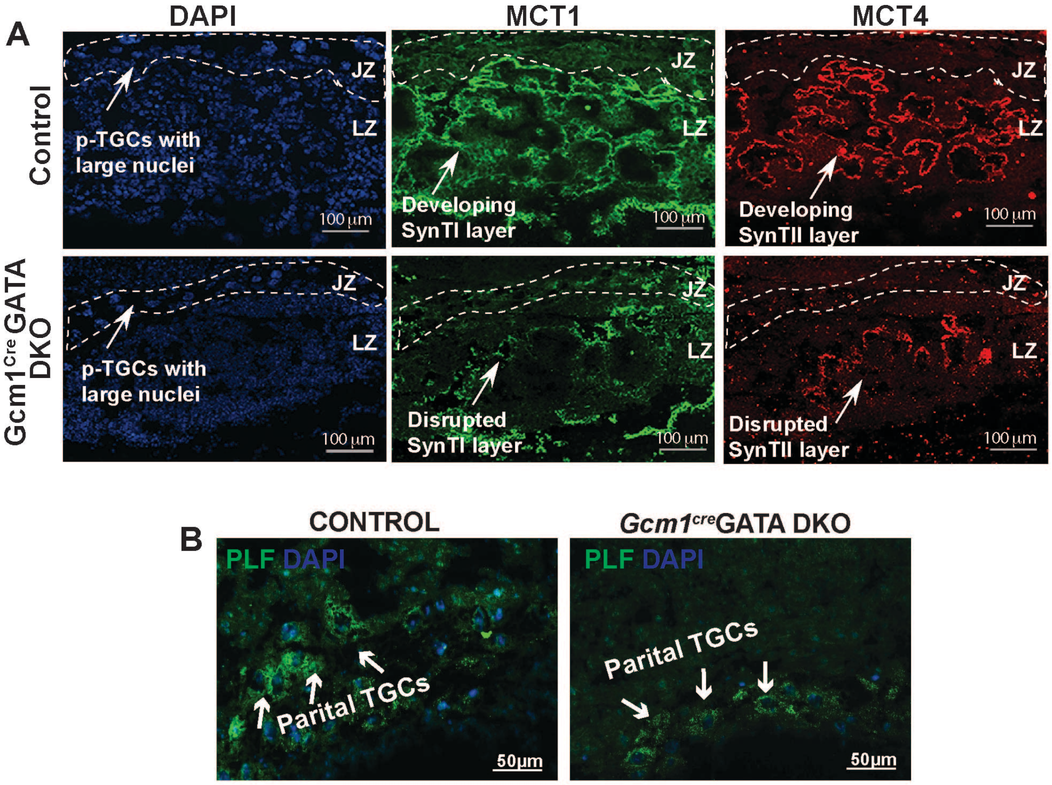

**Figure S2.** (A) Immunofluorescent analysis of the mature SynTI and SynTII layers was done in control and *Gcm1*$^{Cre}$GATA DKO placental labyrinth with MCT1 (green, marks SynTI layer) and MCT4 (red, marks SynTII layer) antibodies. The developing *Gcm1*$^{Cre}$GATA DKO placental labyrinth zone (LZ) lacked mature SynTI and SynTI layers, they were disrupted as compared to juxtaposed layered organization observed in the control (Scale bars, 100 µm.). B, Immunofluorescent analysis of the pTGCs was done in control and *Gcm1* $^{Cre}$GATA DKO placental labyrinth with PL1 (red, marks pTGCs) antibody. The junctional zone (JZ) in both the control and the *Gcm1* $^{Cre}$GATA DKO contained parietal TGCs (p-TGCs). (Scale bars, 50 µm.).

### Fig. S3

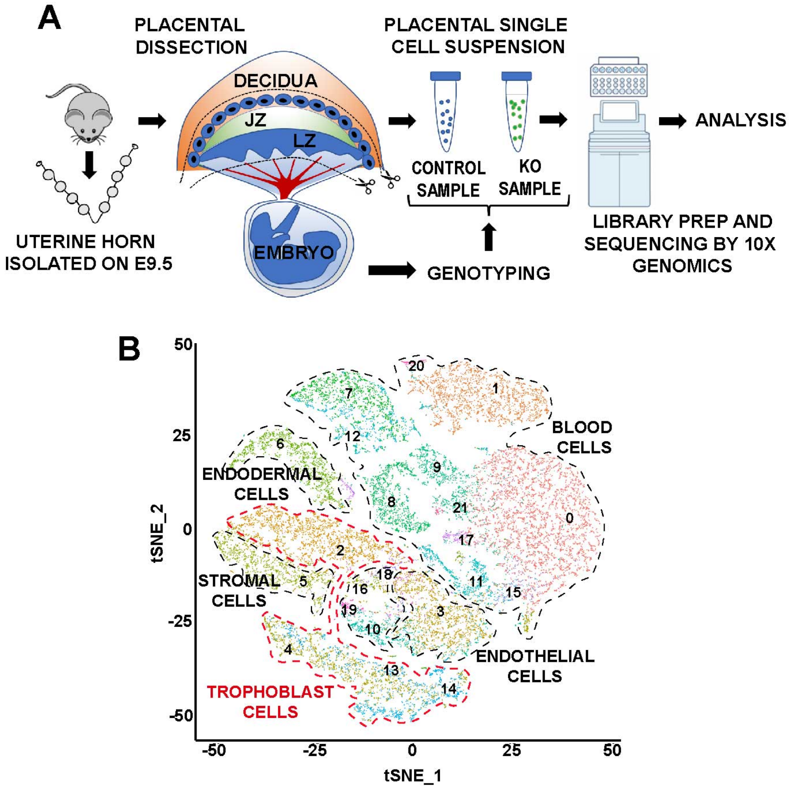

**Figure S3.** (A) Schematic showing the strategy of performing single cell RNA seq (scRNA-Seq) with mouse placenta. The uterine horn at day 9.5 post fertilization was isolated from pregnant mice (cross mentioned in fig 1A). Cartoon representation of the main regions of the placenta – Labyrinth (LZ), Junctional Zone (JZ), and Decidua are shown. Removal of the decidual stroma and the allantois is marked by scissors and cut lines. After removal of the decidua and the embryo proper, the placentae (5 control placentae and 6 *Gcm1*$^{Cre}$ GATA DKO placentae were pooled together to generate the control and knockout samples respectively) were used to make a single cell suspension for sequencing using the 10x genomics platform; the embryo proper was used for genotyping to segregate control and *Gcm1*$^{Cre}$ GATA DKO placental samples. (B) Visualization of the scRNA-Seq data with the analysis plotted in two dimensions by transcriptome similarity using t-distributed stochastic neighbor embedding (t-SNE). t-SNE plot represents all the different cell types observed in the placenta at E9.5, clustered, and plotted according to transcriptome similarity. Clusters were annotated according to canonical marker genes. Each dot represents one cell colored according to assignment by clustering analysis . Dotted lines encircle clusters with common properties segregated into five broad groups.

Fig. S4

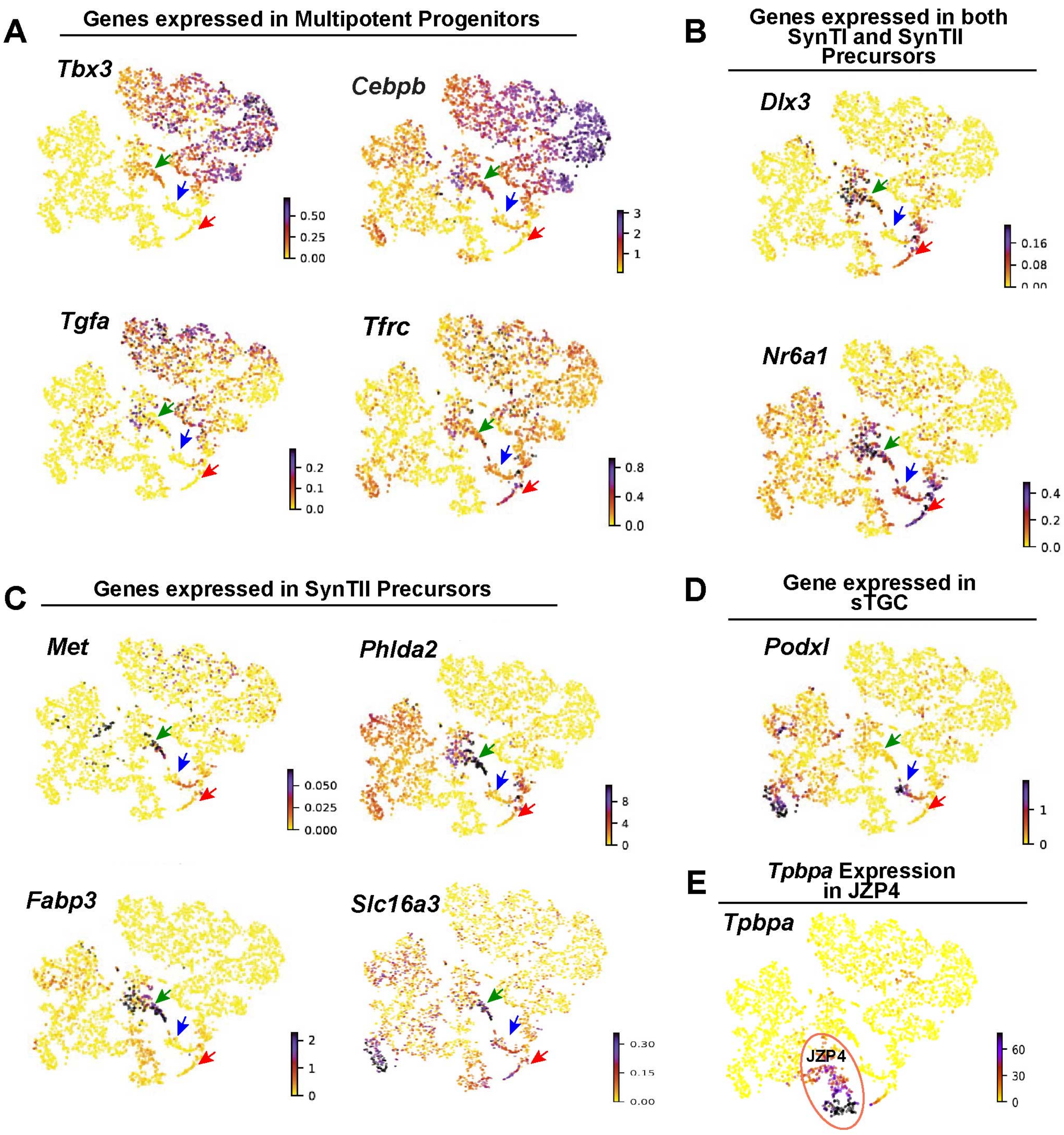

**Figure S4**. RNA velocity plots showing the expression of marker genes in specific clusters, the scale represents the level of expression- yellow (low), red (intermediate) and blue (high). The colored arrows mark specific populations- green=SynTII precursors, blue= S-TGC precursors and red= SynTI precursors. (A) *Tbx3, Cebpb, Tfrc*, Tgfa are highly expressed in the multipotent progenitor cluster. (B) *Dlx3* and *Nr6a* are highly expressed in the SynTI precursors (red arrow), SynTII precursors (green arrow) but absent in S-TGC precursors (blue arrow). (C) *Met, Phlda2, Fabp3* and *Slc16a3* are highly expressed in the SynTII precursors. (D) *Podxl* is expressed by the S-TGC precursors (red arrow). (E) Tpbpa expression is observed only in JZP4 cluster.

Fig. S5

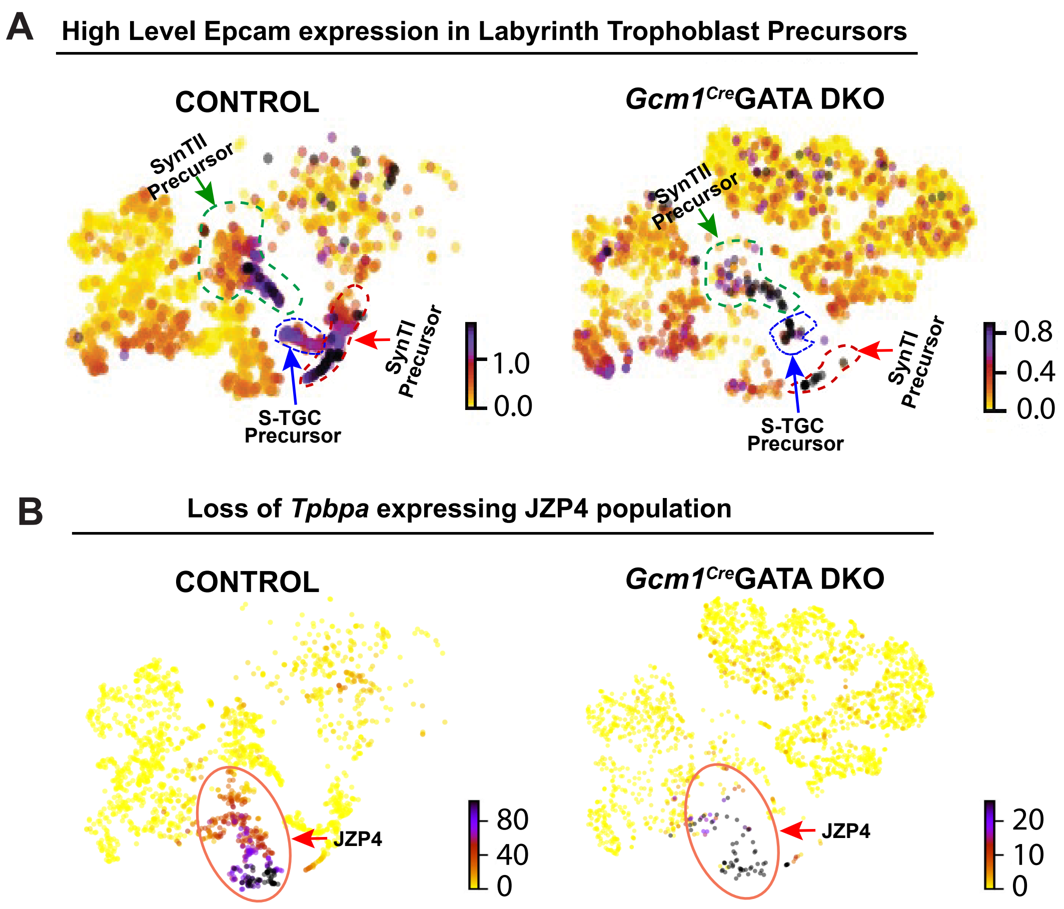

**Figure S5**. RNA velocity plots showing the expression of marker genes in specific clusters, the scale represents the level of expression- yellow (low), red (intermediate) and blue (high). The colored arrows mark specific populations- green=SynTII precursors, blue= S-TGC precursors and red= SynTI precursors. (top) Epcam (High) expression is observed in the SynTI precursors, SynTII precursors and S-TGC precursors in both control as well as *Gcm1$^{Cre}$* GATA DKO placentae. (bottom) Tpbpa expression is observed only in JZP4 cluster in the control but diminishes in the *Gcm1$^{Cre}$*GATA DKO.

Fig. S6

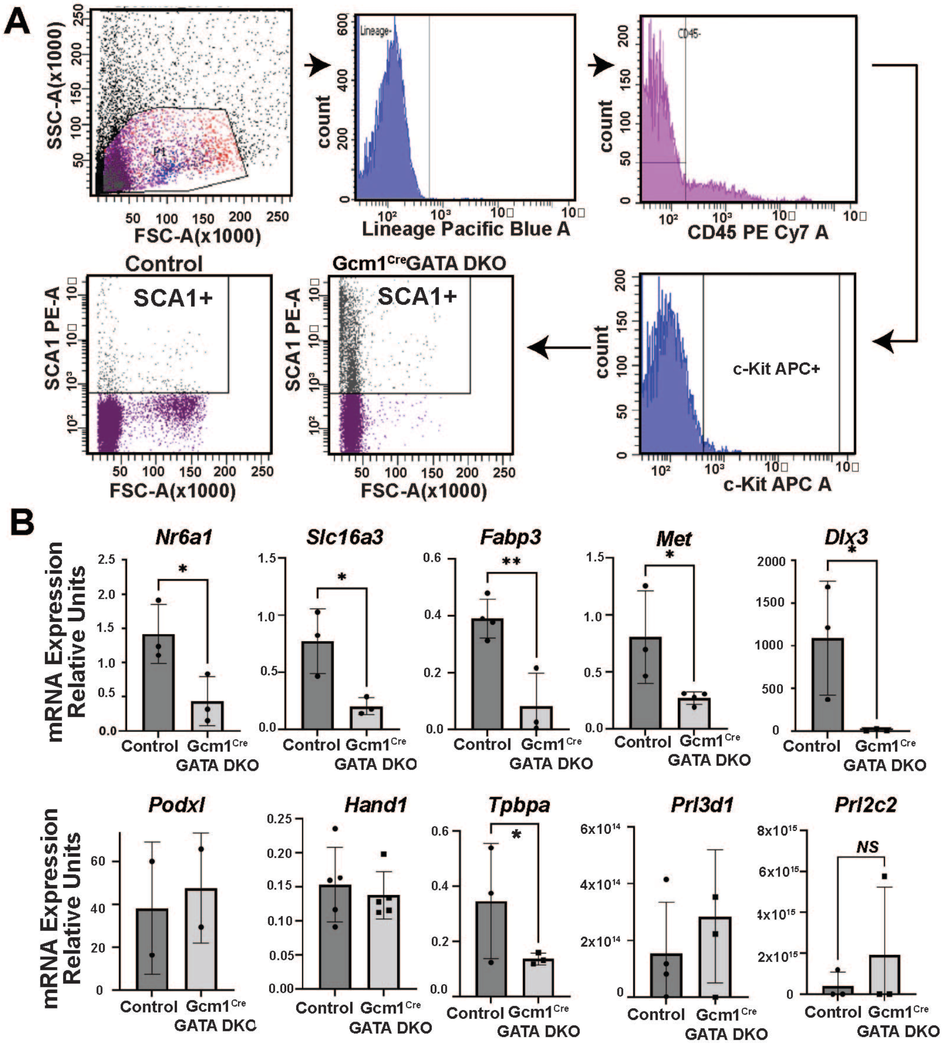

**Figure S6**. (A) FACS sorting of SCA1 (Ly6a) positive trophoblast cells from control and *Gcm1^Cre*GATA DKO placenta (n=3) confirmed the accumulation of these cells in the absence of the GATA factors. Single cell preparation from control and *Gcm1^Cre*GATA-DKO placentae were prepared followed by removal of all the hematopoietic and endothelial cells (Lineage negative, CD34 negative, CD45 negative), and tested percentage of cells that were SCA-1 positive. (B) Quantitative RT-PCR analyses (mean ± SE; n = 3, P ≤ 0.001) reveal downregulation of SynT precursor specific markers like Nr6a1, Slc16a3, Fabp3, Met and Dlx3 in *Gcm1^Cre*GATA DKO placentae compared to control at E9.5. While markers specific for S-TGC precursors like Hand1 and Podxl and junctional zone trophoblasts markers like Tpbpa, Prl3d1 and Prl2c2 remain unchanged.

**Fig. S7**

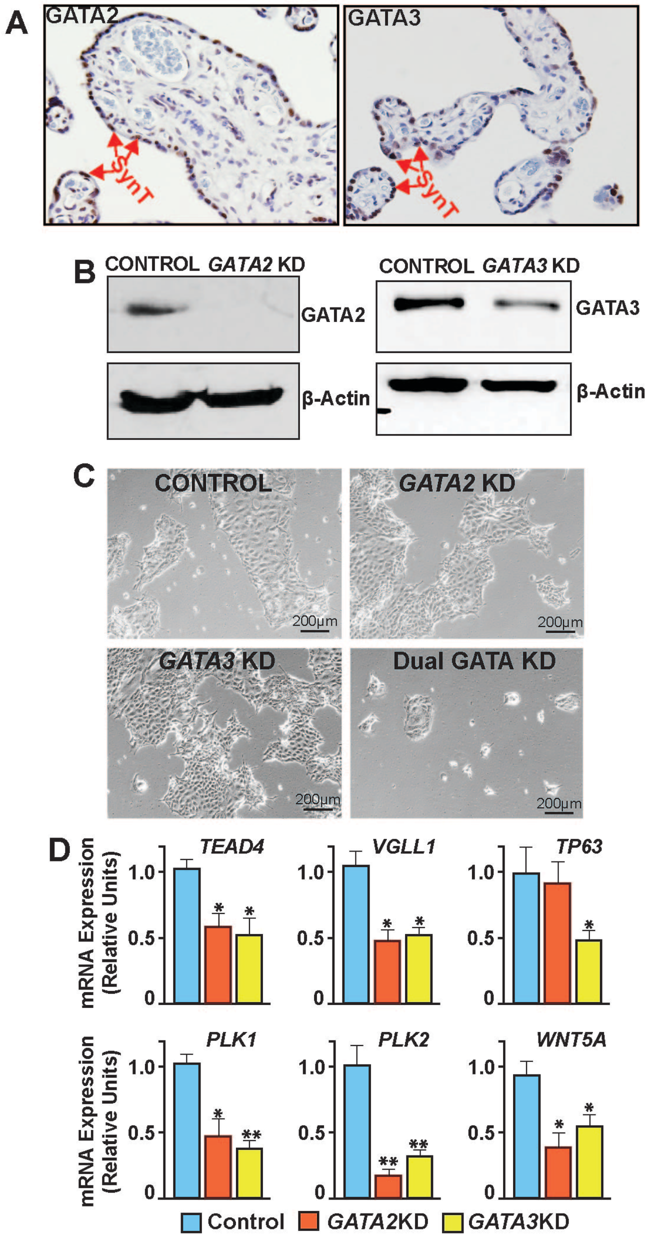

**Figure S7**. (A) Immunohistochemistry showing expression of GATA2 and GATA3 in term (35wk) human placenta. (B) Westernblot analyses showing shRNA-mediated depletion of GATA2 and GATA3 in human TSCs (C) The stem state colony formation and proliferation of scrambled, *GATA2*KD and *GATA3*KD human TSCs was not severely affected after multiple passages (images represent passage 4, P4). But the dual knockdown of GATA2 and GATA3 in human TSCs affected stem state colony maintenance at P0 and could not be passaged further.(D) Quantitative RT-PCR analyses (mean ± SE; n = 3, *,$P \leq 0.05$, **,$P \leq 0.001$) reveal impaired expression of genes, which are important to maintain Human TSC stem-state and proliferation, in *GATA2*KD and *GATA3*KD human TSCs.

**Fig. S8**

**A**

| GENE | Altered in PE/FGR | GATA2 Target | GATA3 Target |
|---|---|---|---|
| *DIO2* | PE/FGR | Yes | Yes |
| *ENG* | PE | No | Yes |
| *FLT1* | PE | Yes | Yes |
| *FSTL3* | PE/FGR | No | Yes |
| *HTRA4* | PE | No | Yes |
| *LEP* | PE | No | Yes |
| *TREM1* | PE/FGR | Yes | No |

**B**

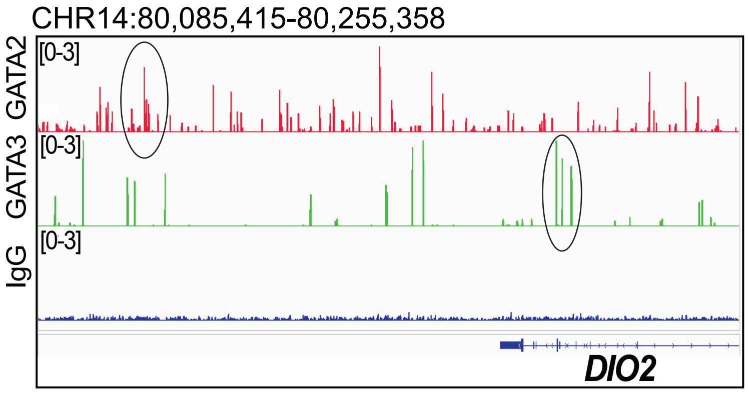

**C**

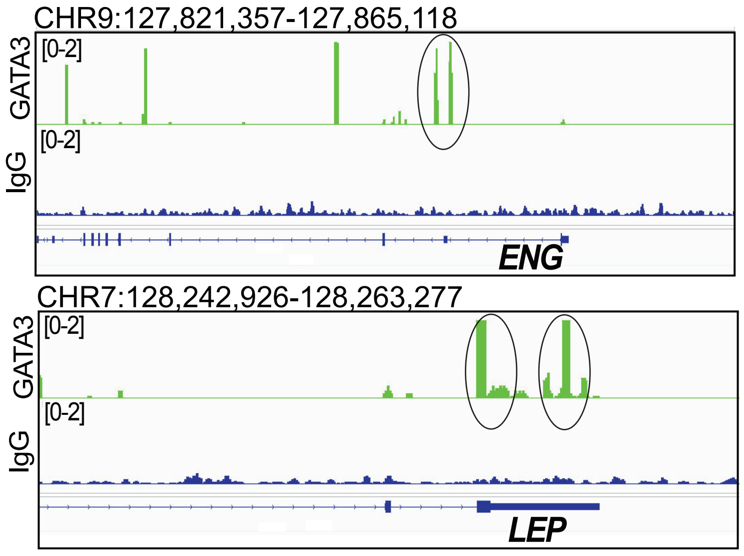

**Figure S8.** (A) The table shows GATA2 and GATA3 target genes (in human TSCs), which are also significantly altered in the placent from pregnancies complicated with PE and/or FGR. These genes are also differentially regulated in *GATA2*KD and *GATA3*KD human TSCs. (B) IGV tracks showing GATA2 and GATA3 binding peaks at the *DIO2* locus. (C) IGV tracks showing GATA3 binding peaks at the *ENG* and *LEP* loci.
